# The *Mycobacterium tuberculosis* Ku C-terminus orchestrates LigD activity through domain-specific interactions

**DOI:** 10.64898/2025.12.12.693941

**Authors:** Dana J. Sowa, Jinfeng Huang, Anmol Marwaha, Asha Subramaniam, Caitlin Doubleday, Monica M. Warner, Jung Ah Byun, Giuseppe Melacini, Sara N. Andres

**Affiliations:** Department of Biochemistry and Biomedical Sciences, McMaster University, Hamilton, Ontario, L8S 4K1, Canada; Michael G. DeGroote Institute for Infectious Disease Research, McMaster University, Hamilton, Ontario, L8S 4L8, Canada; Department of Chemistry and Chemical Biology, McMaster University, Hamilton, Ontario, L8S 4K1, Canada

## Abstract

Bacterial non-homologous end joining (NHEJ) is a DNA double-strand break (DSB) repair pathway that relies on the Ku–LigD complex to alleviate genomic instability. The *Mycobacterium tuberculosis* Ku C-terminus has been highlighted for its role in LigD recruitment to DNA DSBs and stimulation of ligase activity. However, it remains unclear how the Ku C-terminus interacts with and potentially influences other LigD activities. Here, we combine NMR spectroscopy, structural modelling and mutational analysis to define the interaction interface between the Ku C-terminus and LigD. We identify critical residues in Ku (E246, V248, S258, K260, and N266) and the LigD polymerase (D162, V194, R198) and ligase (D522, K579, L580) domains that mediate this interaction. Functional assays reveal that Ku stimulates LigD ligase activity through contacts with both polymerase and ligase domains, while Ku attenuates template-dependent polymerase activity, contrasting previous studies with *Pseudomonas aeruginosa* homologs. Disrupting the Ku–LigD interface, either through Ku or LigD mutations, abolishes ligase stimulation and restores polymerase activity, highlighting a dual regulatory mechanism. Our data supports a model where Ku’s C-terminal region forms a bipartite interface with LigD to balance repair activity. These findings provide mechanistic insight into the Ku–LigD repair mechanism and uncovers species-specific differences in bacterial NHEJ.

**Graphical Abstract:** 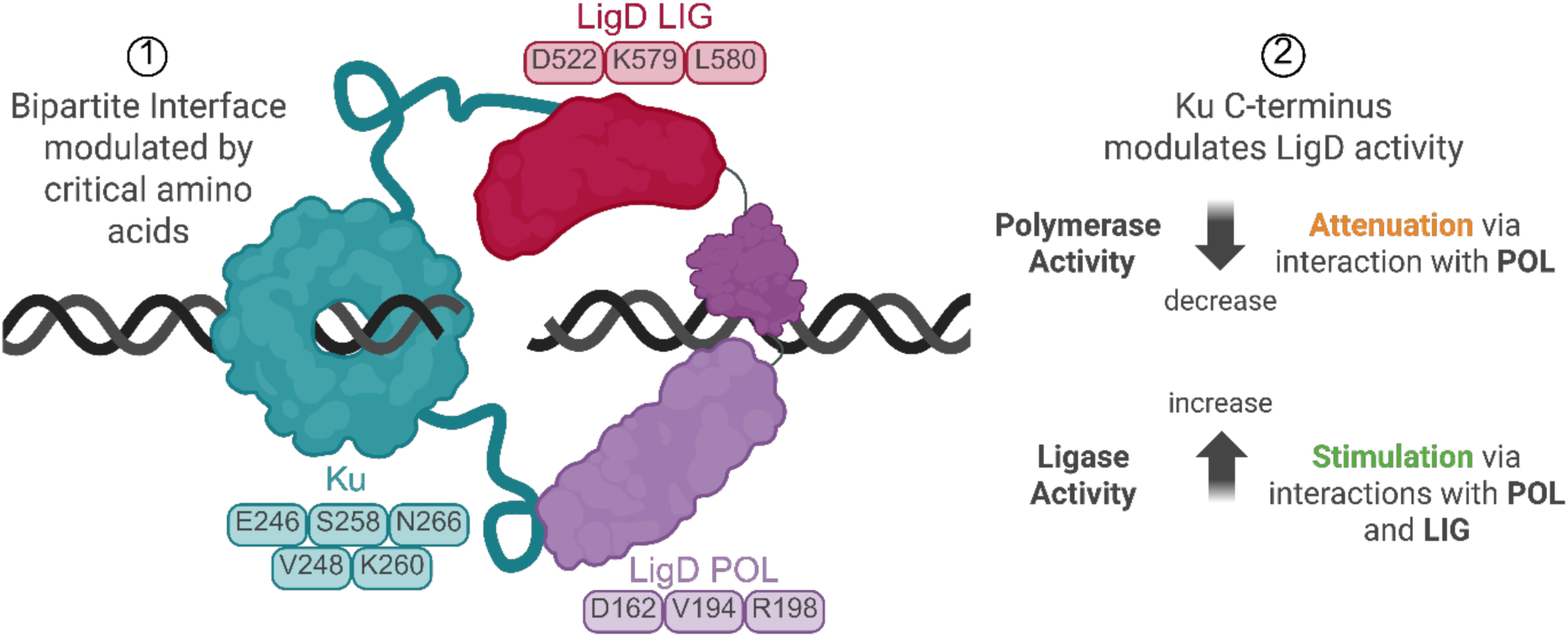

## Introduction

Bacterial genomes are constantly exposed to environmental pressures that cause DNA damage, including double-strand breaks (DSBs)—the most lethal form of DNA damage (1, 2). Bacteria can use non-homologous end joining (NHEJ) repair to mitigate this damage (1, 2), through Ku, a DNA end-binding protein, and LigD, an ATP-dependent ligase with additional enzymatic functions (3). Ku binds and stabilizes DNA ends and recruits its functional partner, LigD, to carry out the repair (1, 4).

Bacterial Ku contains an N-terminal core domain, homologous to the eukaryotic Ku70/Ku80 core domain, which binds DNA through its central pore, and a unique C-terminal region absent in eukaryotic counterparts (3, 5). This C-terminal region can be subdivided into a minimal conserved segment directly following the core, and a highly variable extended region that differs in length and amino acid composition across bacterial species (4, 6, 7). Despite this variability, the Ku C-terminal region plays essential roles in DNA end recognition, bridging, translocation, and serves as the primary binding site for LigD (4, 6, 7).

LigD is unique among ligases as it is a multifunctional enzyme comprised of three domains: an ATP-dependent ligase domain, a polymerase domain, and a 3′-phosphoesterase domain (4, 8). The ligase domain seals nicks in the phosphodiester backbone, while the phosphoesterase domain processes atypical 3′-phosphate DNA ends into 3′-hydroxyl groups for ligation and can also resect 3′ overhangs at DSBs (9–11). The polymerase domain carries out templated synthesis using deoxyribonucleotides (dNTPs) and, preferentially, ribonucleotides (rNTPs), and can add on single-nucleotide, non-templated additions at 5′ overhangs (12, 13).

The enzymatic domains of LigD vary in their composition and arrangement across bacterial species **(Figure 1)**. For example, *Bacillus subtilis* LigD lacks the phosphoesterase domain, acting as a bifunctional enzyme (4, 7), while *Pseudomonas aeruginosa* LigD’s domain arrangement is shifted relative to *Mycobacterium tuberculosis* LigD. Such structural differences likely influence how Ku and LigD interact and how Ku affects LigD activity across species. Notably, the Ku C-terminal region has been identified as the binding hub for the LigD polymerase domain in both *M. tuberculosis* and *B. subtilis*, emphasizing the polymerase domain’s central role in forming the NHEJ repair complex (6, 7).

**Figure 1:**
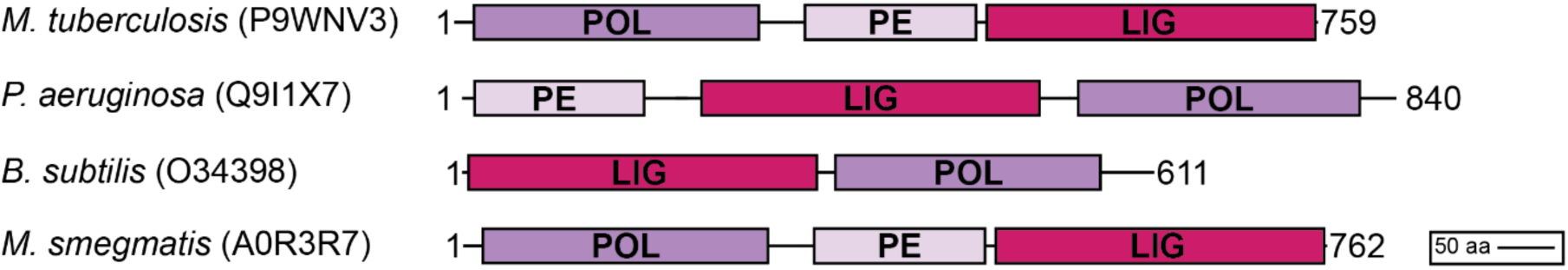
LigD domain arrangement varies across bacterial species. LigD protein domain alignment from model bacterial species *M. tuberculosis* (UniProtKB: P9WNV3), *P. aeruginosa* (UniProtKB: Q9I1X7), *B. subtilis* (UniProtKB: O34398) and *M. smegmatis* (UniProtKB: A0R3R7).

Previous studies have shown that Ku stimulates LigD’s polymerase activity in *P. aeruginosa* (12), and enhances ligase activity in *M. tuberculosis* and *B. subtilis* (6, 7, 14, 15). However, it remains unclear how Ku influences LigD’s polymerase activity in *M. tuberculosis*, and which specific interactions between the Ku C-terminal region and the polymerase domain of LigD are required to form and regulate the Ku–LigD repair complex. Understanding this regulatory interface is key to deciphering species-specific mechanisms of bacterial NHEJ and the balance between repair fidelity and efficiency.

Here, we take a quantitative approach to investigate how disruption of the *M. tuberculosis* Ku–LigD interaction affects LigD function. We find that the Ku C-terminal region forms multiple contacts with the polymerase domain of LigD, which are critical for Ku-stimulated ligase activity. Notably, we show that Ku attenuates LigD template-dependent polymerase activity, differing from the activity observed with homologs from *P. aeruginosa*, and that disrupting these Ku C-terminal interactions restores polymerase function—suggesting that Ku acts as a negative regulator of LigD’s template-dependent polymerase activity. Furthermore, we identify residues in both the polymerase (POL) and ligase (LIG) domains of LigD that interact with the Ku C-terminal region and contribute to the dual regulation of LigD activity. We propose that Ku enhances ligase activity when the Ku C-terminus binds to the polymerase domain. This interaction appears to also repress template-dependent polymerase activity, suggesting Ku regulates LigD ligation and polymerase activities, to coordinate DNA repair.

## Materials and Methods

### Plasmid Cloning

Codon-optimized Ku (UniprotKB: P9WKD9) and LigD (ID: P9WNV3) sequences from *M. tuberculosis* were synthesized by Genewiz and cloned through ligation-independent cloning (LIC)(16), using SspI restriction endonuclease (New England Biolabs, R0132L) and pMCSG7 expression vectors with a TEV-cleavable, N-terminal 6x His-tag (Addgene)(17). Oligonucleotide primers used (Integrated DNA Technologies) are described in **Supplemental Table 1.** All truncations of Ku and LigD were created by LIC using pMCSG7, or by site-directed mutagenesis(18), Ku and LigD point mutations were created using site-directed mutagenesis (18) or Q5 site-directed mutagenesis as per the manufacturer’s instructions (New England Biolabs, E0554S). All plasmids created in this study were verified using whole-plasmid sequencing (Plasmidsaurus).

### Recombinant Protein Expression and Purification

Expression plasmids for all Ku proteins were grown as previously described, with the exception of the following: expression was induced at an OD600 of ∼0.7 with 0.1 mM IPTG for 16 h at 18°C (6). All Ku and LigD proteins were purified as previously described (6). All protein concentrations throughout this paper are expressed as protein concentrations for a monomeric state, including Ku. All purified Ku and Ku mutants had an A260/A280 ≤ 0.7, while all LigD and LigD mutants had an A260/A280 ≤ 0.4, indicating minimal to no DNA contamination. A 12% SDS-PAGE gel visualizing all proteins used in this study can be found in **Supplemental Figure 1**.

### Size-exclusion chromatography coupled to multi-angle light scattering (SEC-MALS)

SEC-MALS for Ku mutants was conducted as previously described (6). All Ku mutants were run on SEC-MALS to ensure oligomeric state was retained as in the wildtype protein.

### Circular dichroism (CD) spectroscopy

500 µL sample volumes of proteins at 0.1 mg/mL were prepared in 50 mM sodium phosphate buffer pH 7.0. Samples were contained in a 0.1 mm quartz cuvette (JASCO). Spectra from 180 nm to 260 nm were collected using Spectra Management Software v.2.10.1.2 (JASCO) and analyzed using Spectra Analysis Software v.2.15.21.1 (JASCO) on a JASCO J-1100 CD Spectrometer. CD spectra collected were subtracted from buffer alone spectra and smoothed using Savitzky-Golay filtering. Data was then analyzed for secondary structure estimation using CD Multivariate Secondary Structure Estimation (SSE) v.2.3.1.1 (JASCO) and compared to a calibration standard created by CD Multivariate Calibration Model Creation v.2.2.1.1 (JASCO). Secondary structure spectra were plotted in Prism v.10.4.1 (GraphPad).

### DNA substrate preparation

DNA substrates used in DNA binding and polymerization assays were purchased as synthetic oligonucleotides from Integrated DNA Technologies (IDT). Oligonucleotide sequences and modifications can be found in **Supplemental Table 2**. Equimolar concentrations of complementary DNA oligonucleotides were resuspended in milliQ water and were annealed at 95°C for 2 minutes and then cooled to 25°C over 45 min. DNA was then purified by ethanol precipitation and resuspended in milliQ water.

### NMR spectroscopy

The NMR samples were prepared in 20 mM sodium phosphate buffer, 150 mM NaCl, pH 6.0 with 3 mm NMR tubes. The Ku C-terminal region peptide (GenScript) was dissolved in the NMR buffer (20 mM sodium phosphate buffer pH 6.0, 150 mM NaCl, 40% D2O) to generate a 500 μM stock solution. The Ku C-terminal peptide encompasses 35 amino acids (239–273) of the *M. tuberculosis* Ku protein (PRLLDEPEDVSDLLAKLEASVKARSKANSNVPTPP). LigD was also exchanged into the NMR buffer to a stock concentration of 43 μM prior to running it through a Superdex 200 Increase 10/300 GL size exclusion column (GE Healthcare) and concentrating it with a 10 kDa Amicon ultra centrifugal filter (Millipore-Sigma). The D2O content was controlled by mixing the 0% D2O SEC elution and the 100% D2O NMR buffer at a 6:4 ratio. NMR experiments were performed on a Bruker Avance 700 MHz spectrometer equipped with a TCI cryoprobe and were acquired at 283 K. To assign the ^1^H NMR spectrum of the Ku peptide, 2D homonuclear experiments (*i.e.* nuclear overhauser enhancement spectroscopy (NOESY) and total correlation spectroscopy (TOCSY) from Bruker pulse programs noesygpph19 and dipsi2gpph19, respectively) were implemented for 500 μM Ku peptide samples as in **Supplemental Figure 2A**. The NOESY spectrum was acquired with a mixing time of 300 ms. The 2D spectra were recorded with 512 t1 increments and 2K data points for the t2 dimension with a spectral width of 7143 Hz and an inter-scan delay of 1.5 s. TOCSY spectra were acquired with a mixing time of 45 ms, and 512 t1 increments and 2K data points for the t2 dimension with a spectral width of 9612 Hz and an inter-scan delay of 2 s. All spectra were processed using topspin (v.4.0.8) and analyzed with NMRFAM-Sparky. Data were processed with a cosine window function in both F1 and F2 dimensions and zero-filled to 4K and 1K data points in F2 and F1 dimensions, respectively. To assess the interaction between Ku peptide and LigD, TOCSY experiments were employed on 150 μM Ku peptide samples with or without 30 μM LigD, shown in **Supplemental Figure 2B**, and 256 t1 increments and 2K data points for the t2 were employed during acquisition. The HN and Hα cross-peak intensities for each residue were fitted with Gaussian function implemented in NMRFAM-SPARKY (the ambiguous assignments and overlapped peaks are excluded in this work) to evaluate binding interactions, and greater reductions in intensity upon LigD binding to the peptide are assumed to indicate stronger interactions.

### DNA Binding Assays

DNA binding assays were performed for the Ku mutant proteins with 10 nM fluorescein-labelled 40 bp dsDNA as previously described (6, 19). Apparent dissociation constants were obtained through calculation of specific binding with Hill Slope in Prism v.10.4.1 (GraphPad). p-values were calculated by a two-tailed Welch’s t-test in Prism v.10.4.1 (GraphPad).

### Fluorescence Polarization

Fluorescence polarization was used to measure LigD binding affinity to a 50 nM fluorescein-labelled 40 bp dsDNA as previously described (6). Apparent dissociation constants were obtained through calculation of specific binding with Hill slope in Prism v.10.4.1 (GraphPad). *p*-values were calculated by a two-tailed Welch’s *t*-test in Prism v.10.4.1 (GraphPad).

### Microscale Thermophoresis (MST)

The MST experiments were adapted following a previously published protocol (20). Ku and LigD mutant proteins were labeled using the Monolith His-Tag Labeling Kit RED-tris-NTA 2^nd^ generation (NanoTemper Technologies), following manufacturer’s protocol in MST buffer (50 mM HEPES pH 8.0, 50 mM NaCl, 5 mM MgCl2, 0.05% (v/v) Tween-20) at room temperature for 1 hour in the dark. Unreacted dye was removed by centrifugation at 15000 x g for 10 minutes at 4°C. RED-tris-NTA labelled mutant Ku and LigD proteins were diluted to 100 nM with MST buffer, while wild-type Ku and LigD proteins underwent TEV cleavage in S200 buffer (50 mM HEPES pH 8.0, 400 mM NaCl, 10% (v/v) glycerol) to remove the His-tag. Two-fold serial dilutions of wild-type proteins were prepared in MST buffer, producing concentrations ranging from 150 µM to 2.3 nM. Ligand (RED-tris-NTA labelled proteins) and analytes (TEV-cleaved proteins) were incubated at room temperature for 20 minutes. Standard capillaries’ (NanoTemper Technologies) were used for all experiments with Ku mutants while premium capillaries (NanoTemper Technologies) were used for all LigD mutant experiments. Upon loading of samples into the capillaries, binding affinities were measured at 25°C. Data was collected using M.O. control software v.2.6.2 (NanoTemper Technologies) and fitting curves and KD values were calculated from 3 technical replicates using Prism v.10.4.1 (GraphPad), using the one-site specific binding equation. *p*-values were calculated by a two-tailed Welch’s *t*-test in Prism v.10.4.1 (GraphPad).

### Template-dependent DNA Polymerase Assay

DNA polymerase assays were performed for the Ku wildtype and mutant proteins and the LigD wildtype mutant proteins using an assay adapted from Zhu *et al.* (2010) (12). 20 µL reaction mixtures containing polymerase buffer (50 mM HEPES pH 7.5, 5 mM MgCl2, 5mM DTT), 25 µM of each dNTP (dATP, dTTP, dCTP and dGTP) and 40 nM of a fluorescein-labelled 18-bp DNA substrate with an 18-nucleotide (nt) overhang were incubated with 0.5 µM LigD in either the presence or absence of 1 µM Ku at 37°C for 5 minutes. Samples containing Ku were allowed to pre-incubate with the reaction buffer and DNA for 10 minutes prior to adding LigD to allow Ku to interact with DNA. Reactions were quenched at 0, 0.5, 1, 1.5, 2, 3, 4 and 5 min, with 20 μL of polymerase quench buffer (98% formamide, 10 mM EDTA pH 8.0), and denatured at 95°C for 10 minutes prior to resolving on a 20% denaturing PAGE gel. Electrophoresis was performed in 1X Tris-Borate-EDTA buffer at 200V for 2 hours to resolve the samples. Gels were imaged using an Amersham Typhoon (GE Healthcare) and analyzed using ImageJ. Experiments were performed in n=3 technical replicates, and rates of polymerization were calculated using simple linear regression using Prism v.10.4.1 (GraphPad). *p*-values were calculated by a two-tailed Welch’s *t*-test in Prism v.10.4.1 (GraphPad).

### DNA Ligation assay

DNA ligation assays were performed for the Ku wildtype and mutant proteins and LigD wildtype and mutant proteins as previously described, using pUC-19 linearized by KpnI, leaving behind sticky DNA ends (6). DNA ligation results were plotted as a function of the phosphate released (nmol versus time (min)). Phosphate released was calculated from a standard curve (6). Slopes from these plots were calculated by simple linear regression in Prism v.10.4.1 (GraphPad), which corresponded to the rate of the reaction. *p*-values were calculated by a two-tailed Welch’s *t*-test in Prism v.10.4.1 (GraphPad).

## Results

### Conserved amino acids of the Ku C-terminus are important for the Ku-LigD interaction

Previous biochemical studies by us and others have shown that the Ku C-terminus binds to the polymerase domain of LigD (6, 7), for both *B. subtilis* and *M. tuberculosis* homologs, but the critical residues mediating this interaction are unknown. By using a synthetic peptide of the *M. tuberculosis* Ku C-terminus with the LigD polymerase domain, we acquired 2D NOESY and TOCSY spectra to identify conformational changes of the Ku C-terminal peptide when interacting with the LigD polymerase domain. A loss of signal intensity at a specific amino acid position of the Ku C-terminal peptide when bound to the LigD polymerase domain (∼31 kDa), compared to the peptide alone, indicates a potential interaction between the two proteins. From this experiment, we identified ten amino acids that lose intensity in the presence of the LigD polymerase domain **(Figure 2A, Supplemental Figure 2)**. Of these ten amino acids, E246, V248, S258, V259, K260, A261, R262 and S263 are primarily conserved amongst bacterial species, while A265 and N266 are not.

**Figure 2:**
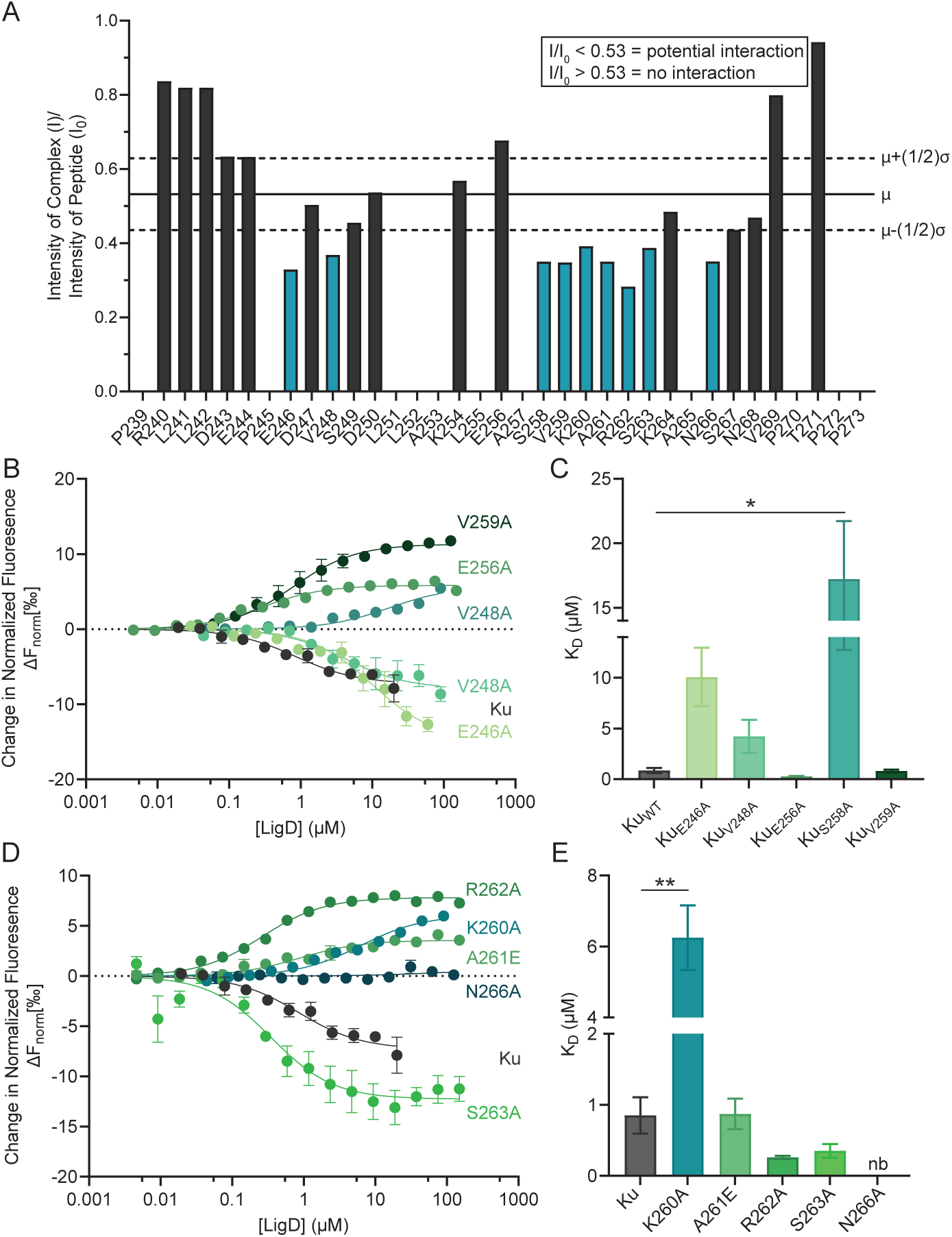
The Ku C-terminus contains important contacts for the Ku-LigD interaction. (A) NMR intensity ratios of the *M. tuberculosis* LigD polymerase domain and a peptide of the Ku C-terminus vs. the peptide alone. Highlighted in teal are the 10 residues of the Ku C-terminal region in the presence of LigD that fall below the (I)/(Io) average of 0.53 (µ). Average ± ½ the standard deviation (σ) is represented by the dashed lines. (B) Binding curves of the Ku-LigD interaction with Ku minimal C-terminal region mutants, measuring the change in normalized fluorescence (ΔFnorm) of the Ku ligand over the concentration of LigD analyte (µM). Data is fitted with non-linear regression using the binding saturation – one site, specific function in Prism v.10.1.4 (GraphPad). (C) Apparent dissociation constants (KD) for LigD binding Ku and Ku mutants from (B). (D) Binding curves of the Ku-LigD interaction, with Ku C-terminal region mutants, measuring the change in normalized fluorescence (ΔFnorm) of the Ku ligand over the concentration of LigD analyte (µM). Data is fitted with non-linear regression using the binding saturation – one site, specific function in Prism v.10.1.4 (GraphPad). (E) Apparent dissociation constants (KD) for LigD binding Ku and Ku mutants in (D). Data are plotted as the mean ± standard error measure for n=3 technical replicates. **p<0.1, *p<0.05 (two-tailed t-test). Ku MST values are plotted twice to allow comparison with Ku mutants. MST raw traces are available in Supplemental Figure 5.

The NMR spectral intensity change could indicate that there is a direct Ku-LigD interaction and/or a conformational change in the Ku peptide that is triggered upon binding LigD. Therefore, we created single-point mutants of the 10 amino acids in the wild-type Ku protein that were identified through NMR to understand their impact on the Ku-LigD interaction. We verified that all Ku mutants had retained proper secondary structure folding and a homodimeric state using CD Spectroscopy and SEC-MALS **(Supplemental Table 3, Supplemental Figure 3-4)**. We then used microscale thermophoresis (MST) to measure changes in binding affinity of the Ku mutants to wild-type LigD **(Figure 2 B-E, Supplemental Table 4, Supplemental Figure 5)**. Using MST, the binding affinity between Ku and LigD was in the µM range (KD = 0.85±0.26 µM). Mutant Ku proteins E246A, V248A, S258A, and K260A had 5 to 20-fold decreased binding affinity, suggesting that these amino acids contribute to the Ku-LigD interaction. Mutant Ku proteins E256A, R262A, and S263A had slightly increased binding affinity (2.5-3-fold), suggesting that these amino acid side chains are not critical to the Ku-LigD interaction. Lastly, Ku mutant N266A resulted in a total loss of binding to LigD, suggesting that N266 is essential for the interaction of Ku and LigD.

### Interactions with the Ku C-terminus are critical for stimulating ligase function

Previously, we proposed that Ku may regulate LigD’s enzymatic activity via an allosteric mechanism by interacting with the polymerase domain to elicit an effect in the ligase domain (6). Here, we investigated whether specific amino acids in the Ku C-terminal region that interact with the polymerase domain of LigD also impair ligase activity, supporting the proposed allosteric mechanism. We hypothesized that if these mutations weakened the Ku-LigD interaction, the corresponding Ku mutants would fail to stimulate ligase activity.

We found that ligation rates in the presence of Ku mutants E246A, V248A and N266A, were comparable to that of LigD alone **(Figure 3, Supplemental Table 5)**. These results support our hypothesis that a loss of interaction between Ku and LigD would result in loss of stimulation, indicating these amino acids are critical for Ku-stimulated ligase activity. However, in the presence of Ku K260A the ligation rate increased 1.5-fold compared to LigD alone but was 1.3-fold slower compared to ligation in the presence of Ku. This result suggests that K260 is also important for the stimulation of LigD ligase activity, but not to the same extent as the other amino acids observed in this study.

**Figure 3:**
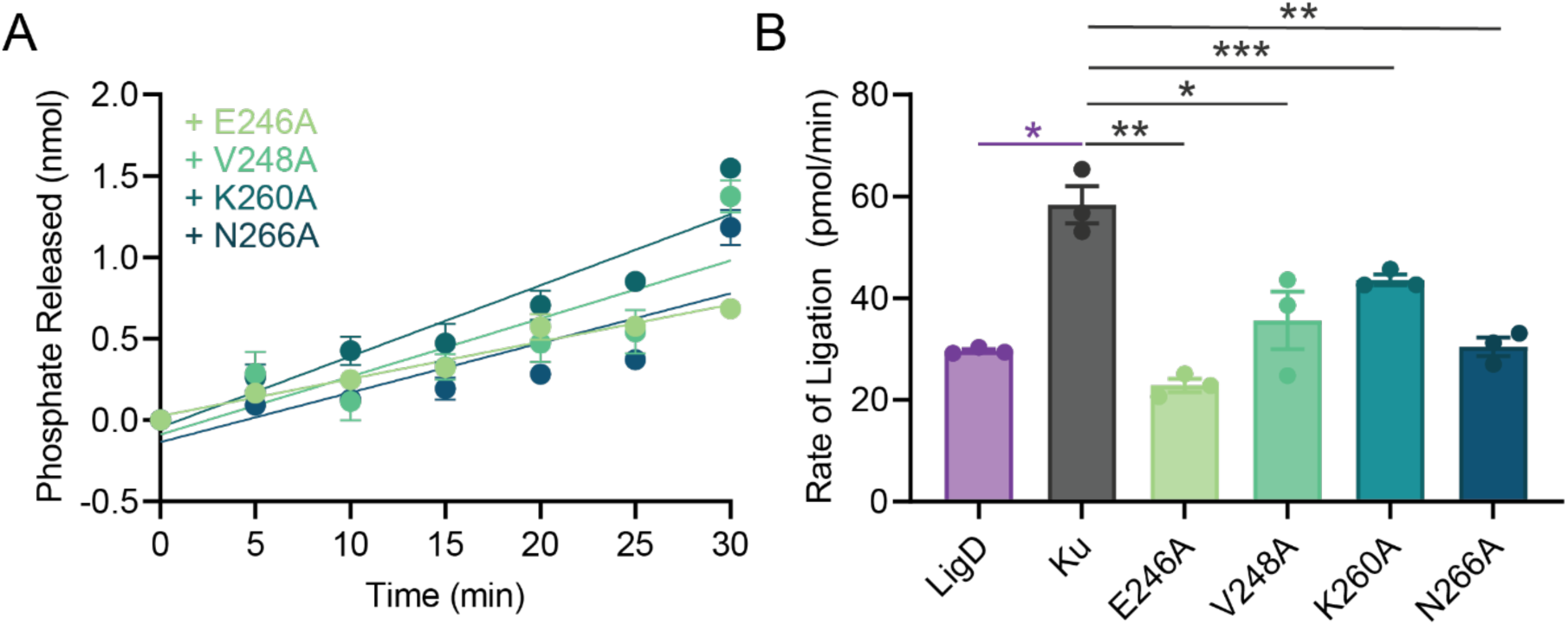
The Ku-LigD interaction is important for efficient repair of DNA. (A) Ligation of the linearized pUC19 plasmid in the presence of Ku C-terminal mutants E246A, V248A, K260A and N266A. Data were plotted as a function of phosphate released (nmol) over time (min). (B) Rates for ligation of linearized pUC19 plasmid in the presence of Ku C-terminal mutants E246A, V248A, K260A and N266A. All data plotted are the mean ± standard error measure. n=3 technical replicates. ***p<0.001, **p<0.01, *p<0.05 (two-tailed t-test) in comparison to LigD (purple lines) or Ku +LigD (grey lines). Ligation rates of LigD and Ku+LigD in (B) are replotted here from (6) to allow for comparison between Ku mutants.

### LigD template-dependent polymerase activity is not stimulated by M. tuberculosis Ku

While Ku stimulates LigD ligase activity in both *P. aeruginosa* and *M. tuberculosis,* it remains unclear whether Ku similarly affects LigD template-dependent polymerase activity in *M. tuberculosis*. To address this, we adapted a polymerase assay from Zhu *et al*. (2010) and measured the rate at which *M. tuberculosis* LigD adds dNTPs in a template-dependent manner **(Figure 4, Supplemental Table 6)** (12). The rate of nucleotide addition for LigD alone is 0.94±0.07 nM/min. However, upon the addition of Ku, the rate of addition decreases to 0.70±0.04 nM/min, contrary to what is observed with protein homologs from *P. aeruginosa* (12), indicating that *M. tuberculosis* Ku does not stimulate LigD’s template-dependent polymerase activity.

**Figure 4:**
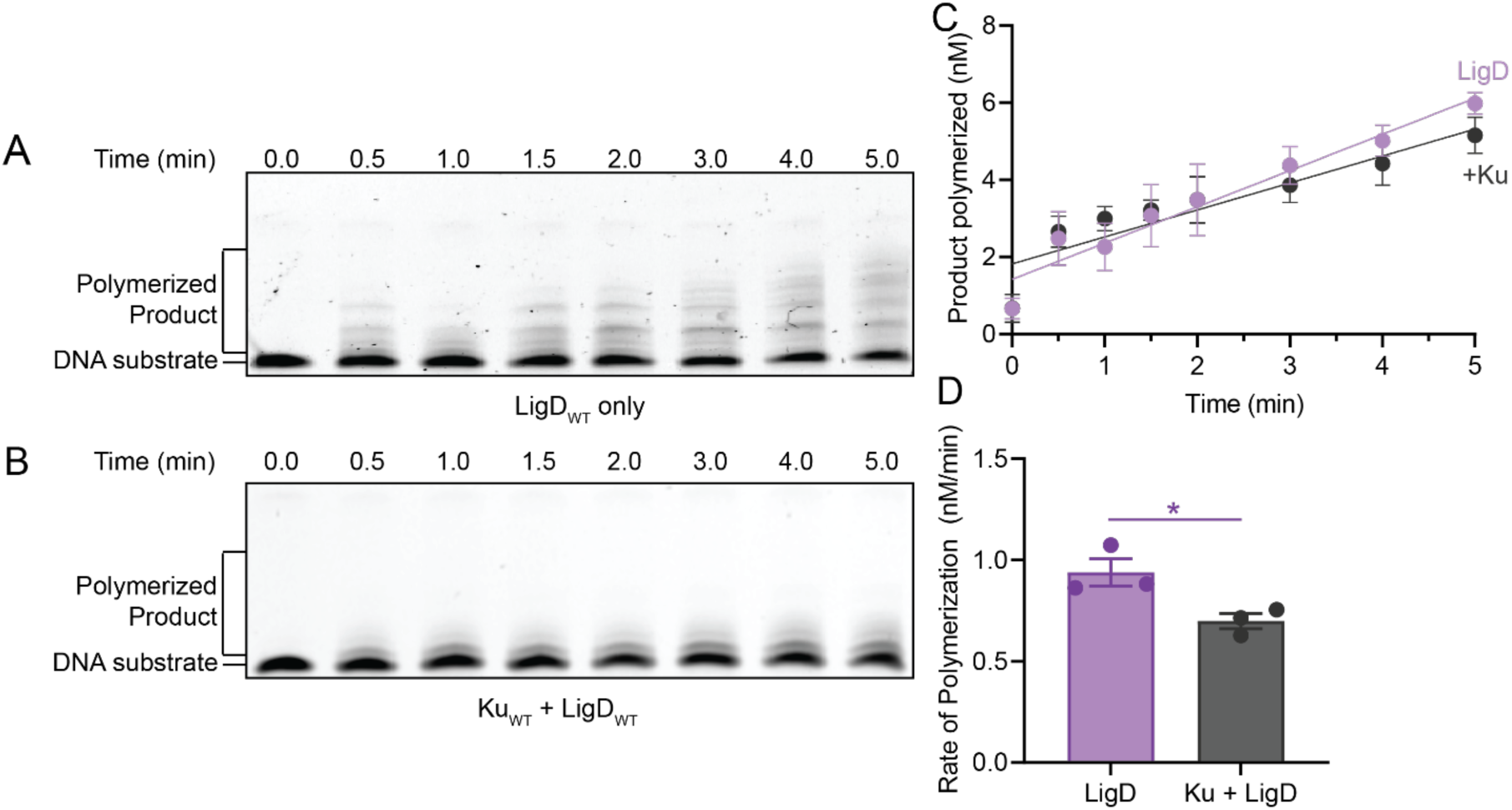
***M. tuberculosis* Ku does not stimulate template-dependent LigD polymerase activity.** The ability of LigD to add dNTPs to an 18-nucleotide overhang templated substrate was visualized on a 20% denaturing-PAGE, for (A) LigD alone, and (B) in the presence of Ku. (C) Polymerization of the 18-nucleotide overhang templated substrate by LigD alone or in the presence of Ku. Data were plotted as a function of product polymerized (nM) over time (min). Slopes were calculated by simple linear regression in Prism v.10.1.4 (GraphPad). (D) Rates for polymerization of 18-nucleotide overhang templated substrate by LigD alone, or in the presence of Ku. All data plotted are the mean ± standard error measure. n=3 technical replicates. *p<0.05 (two-tailed t-test).

We also performed these assays across a range of Ku concentrations to confirm that the observed attenuation of polymerase activity was not concentration dependent. We identified that this attenuation effect was consistent across all concentrations of Ku tested (0.625 – 20 μM), suggesting that Ku does not modulate LigD polymerase activity in a dose-dependent manner, but instead may act as a general attenuator of its activity in *M. tuberculosis* **(Supplemental Figure 6)**.

### The Ku C-terminus is necessary for LigD polymerase activity

NMR and MST studies with the Ku C-terminus identified critical amino acids for the Ku-LigD interaction, therefore we sought to investigate whether disrupting this interaction would restore LigD template-dependent polymerase activity, by relieving any inhibitory effect imposed by Ku. We observed that in the presence of Ku mutants E246A, S258A and N266A, there was no significant change in polymerase activity (1.03±0.05 nM/min, 0.90±0.09 nM/min and 0.85±0.08 nM/min respectively), compared to LigD alone (rate = 0.94±0.07 nM/min) **(Figure 5, Supplemental Table 7, Supplemental Figure 7)**. These results supported our hypothesis, suggesting that disrupting the interaction between Ku and LigD would restore polymerase activity. However, we observed that in the presence of Ku V248A, polymerase activity was maintained at a rate of 0.65±0.11 nM/min, similar to that of LigD in the presence of Ku. Interestingly, in the presence of Ku K260A, polymerase activity was attenuated to a rate of 0.26±0.03 nM/min.

**Figure 5:**
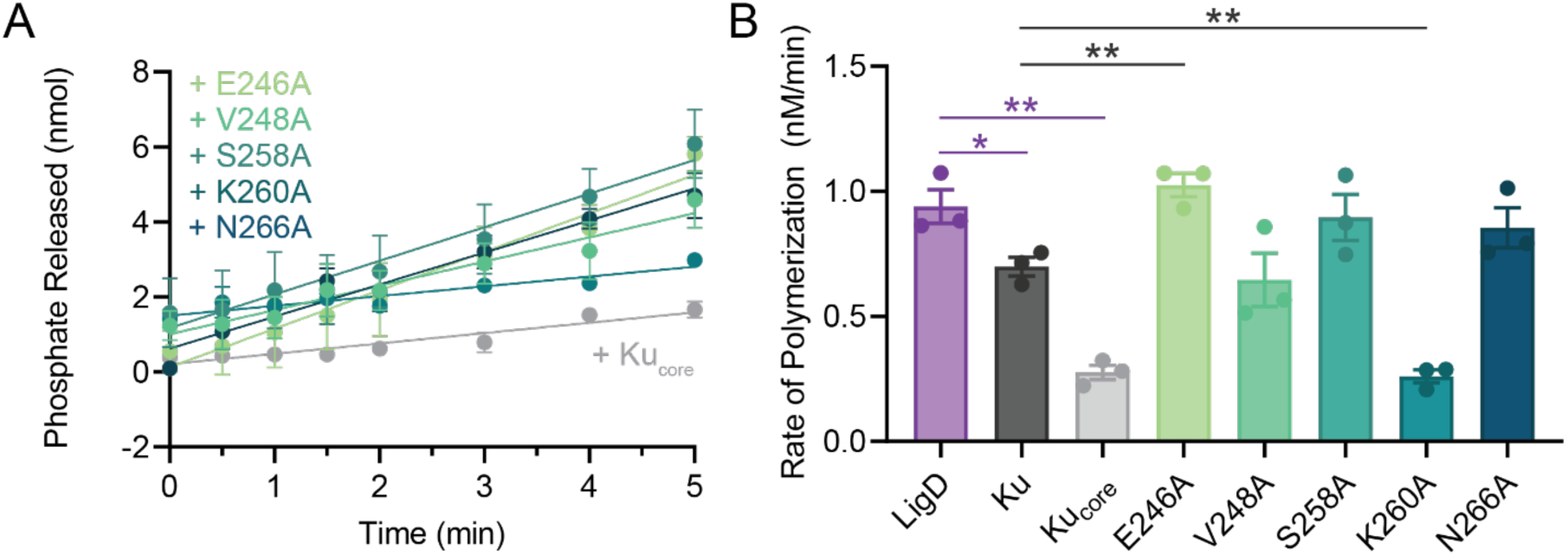
Disruption of the Ku-LigD interaction restores polymerase activity. (A) Polymerization of the 18-nucleotide overhang templated substrate in the presence of Ku C-terminal mutants. Data were plotted as a function of product polymerized (nM) over time (min). Slopes were calculated by simple linear regression in Prism v.10.1.4 (GraphPad). (B) Rates for polymerization of the 18-nucleotide overhang templated substrate in the presence of Ku C-terminal mutants. All data plotted are the mean ± standard error measure. n=3 technical replicates. **p<0.01, *p<0.05 (two-tailed t-test) in comparison to LigD rate (purple lines) or Ku+LigD (grey lines). LigD and Ku+LigD polymerase rates from Figure 4 are replotted here to allow for comparison with Ku mutants. Representative gel images of polymerization data are in Supplemental Figure 7.

In addition to the single point mutations, we investigated LigD polymerase activity in the presence of a Ku mutation which has the entire C-terminus removed (Kucore). Previously, we found that in the presence of Kucore, LigD ligase activity only had slight stimulation of DNA DSB ligation, and nick sealing was abolished (6).Here, we found that Kucore further decreased the rate of LigD polymerization to 0.27±0.01 nM/min, similar to the Ku K260A mutation, showing that removal of the Ku C-terminus impairs LigD DNA polymerization. These results suggest that the Ku C-terminus is necessary for polymerase activity to occur, although this requirement may not solely depend on a Ku-LigD interaction, and that other molecular mechanisms may be involved.

### Ku-DNA interaction is regulated by the Ku C-terminus

We determined that the Ku C-terminus is required for template-dependent polymerase activity, but since Ku attenuates this activity, we wanted to explore if Ku-DNA binding could explain our observations. Therefore, we conducted DNA binding studies for the Ku mutant proteins with a 40-base pair (bp) blunt-ended substrate **(Figure 6, Supplemental Table 8, Supplemental Figure 8)**. In our previous work, we determined that wild-type Ku bound a 40 bp blunt-ended substrate with an affinity of 3.15±0.74 µM, Ku S258A bound DNA with a ∼2-fold tighter affinity, and Kucore bound DNA with an 45-fold higher affinity (6). Here, Ku mutants V248A and N266A had DNA binding affinities similar to that of Ku (V248A, KD = 3.19±0.46 µM; N266A, 3.62±0.52 µM), indicating that these residues do not play a direct role in binding DNA. Ku mutants E246A and K260A both had significantly tighter DNA binding affinities, increasing ∼2-fold to ∼8-fold respectively, compared to Ku, suggesting that E246 and K260, like the Ku C-terminal tail, limit DNA binding affinity. However, the DNA binding activity of Ku and the Ku mutants do not correlate with the observed LigD polymerase activity in the presence of Ku.

**Figure 6:**
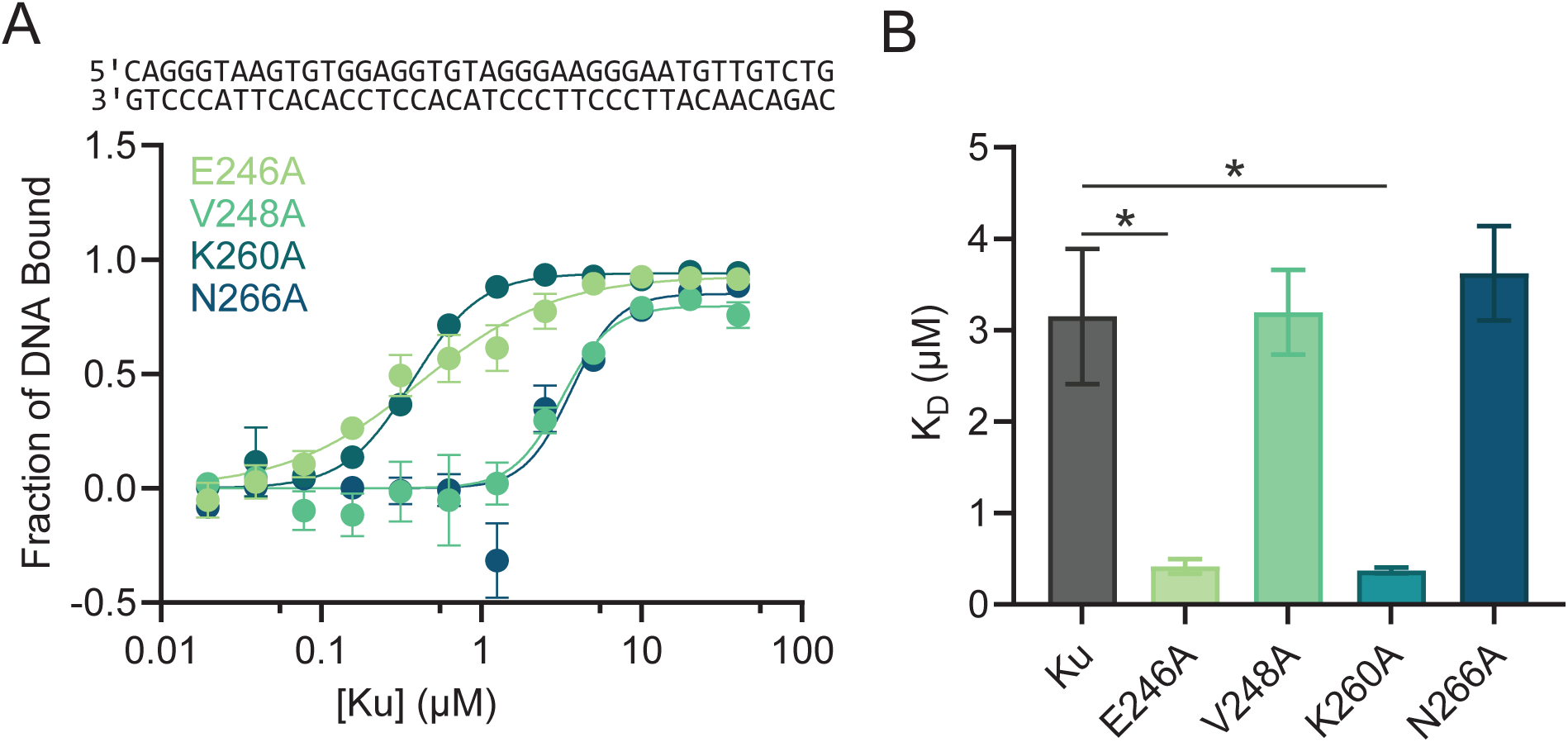
The Ku C-terminal tail regulates DNA binding. (A) DNA binding curves for Ku mutants E246A, V248A, K260A and N266A, binding to a 40 bp DNA substrate. Data is fitted with non-linear regression using the specific binding with hill slope function in Prism v.10.1.4 (GraphPad). (B) Apparent dissociation constants (KD) for Ku and Ku E246A, V248A, K260A and N266A binding to the 40 bp DNA substrate. All data plotted are the mean ± standard error measure. n=3 technical replicates. *p<0.05 (two-tailed t-test). DNA binding apparent dissociation constant of Ku is replotted here from (6) to allow for comparison with Ku mutants. Representative gel images of binding data are in Supplemental Figure 8.

### Comparing the effects of Ku homologs on LigD polymerase activity

Our findings contrast with previous reports that *P. aeruginosa* Ku stimulates LigD template-dependent polymerase activity (12). Therefore, we wanted to directly compare LigD polymerase activity between the homologs. Using the templated addition polymerization assay for *P. aeruginosa* LigD, we observed a polymerization rate of 0.65±0.02 nM/min, which increased 2-fold to 1.29±0.05 nM/min upon addition of Ku **(Figure 7, Supplemental Table 9, Supplemental Figure 9)**. In contrast, the *M. tuberculosis* LigD polymerization rate decreased in the presence of Ku. Notably, *M. tuberculosis* LigD alone displayed a 1.4-fold higher polymerization rate than its *P. aeruginosa* counterpart, but in the presence of Ku, its activity was 1.4-fold lower—highlighting species-specific differences in Ku-mediated regulation of LigD polymerase activity.

**Figure 7:**
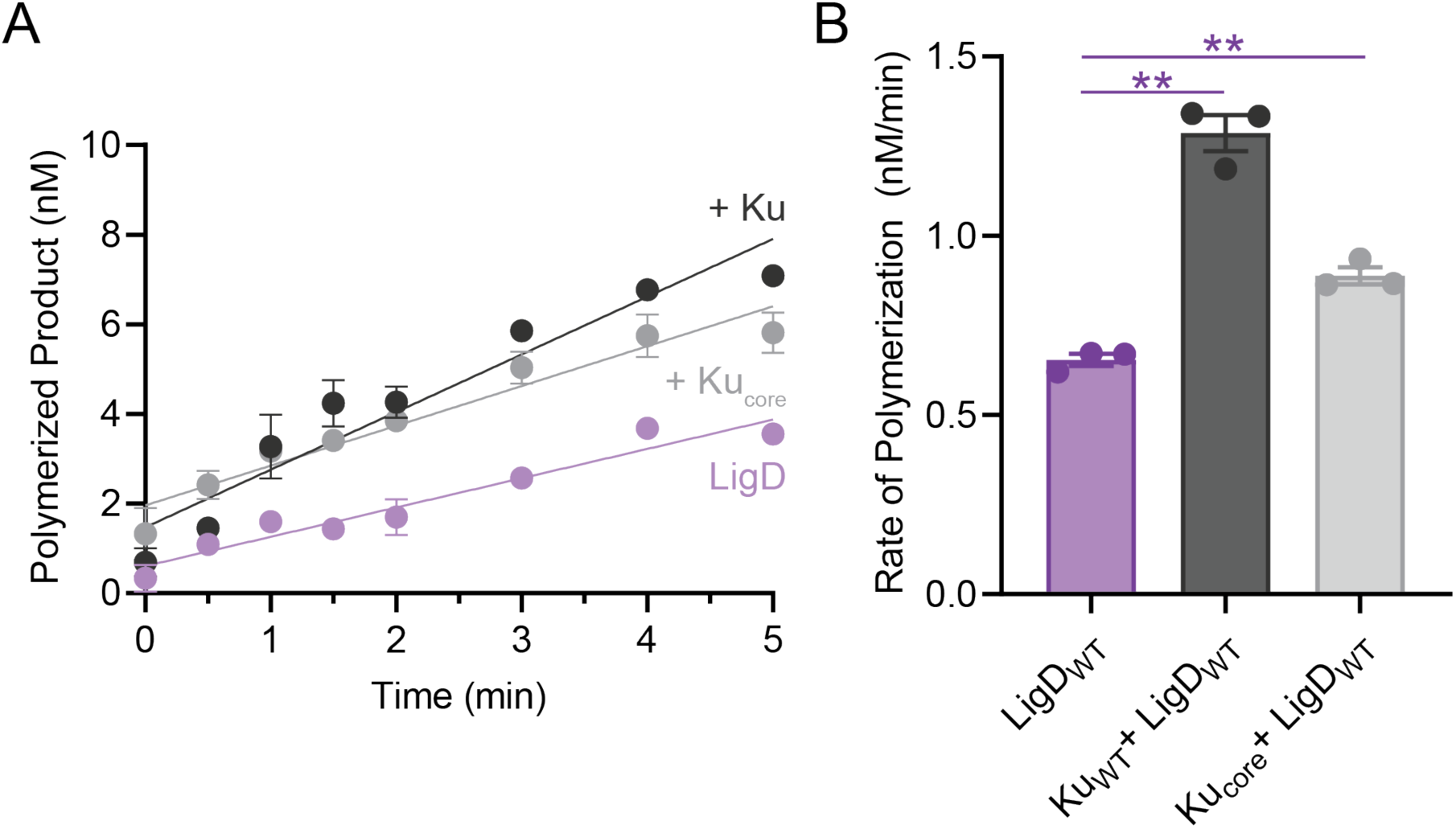
***P. aeruginosa* Ku stimulates LigD template-dependent polymerase activity.** (A) Polymerization of the 18-nucleotide overhang templated substrate by LigD alone or in the presence of Ku or Kucore. Data are plotted as a function of product polymerized (nM) over time (min). Slopes were calculated by simple linear regression in Prism v.10.1.4 (GraphPad). (B) Rates for polymerization of 18-nucleotide overhang templated substrate by LigD alone, or in the presence of Ku or Kucore. All data plotted are the mean ± standard error measure. n=3 technical replicates. **p<0.01, *p<0.05 (two-tailed t-test). Representative gel images of binding data are in Supplemental Figure 9.

In addition, we wanted to observe what would happen to polymerase activity when the *P. aeruginosa* Ku C-terminus was removed. Therefore, we created a mutation for a premature stop codon in *P. aeruginosa* Ku to create a Kucore mutant that removed the entire C-terminal region (amino acids 239-293). We identified that, unlike its counterpart in *M. tuberculosis*, *P. aeruginosa* Kucore did not impair polymerase activity, resulting in a rate of polymerization of 0.89±0.02 nM/min. This 1.3-fold increase in activity compared to LigD alone showed slight stimulation. However, it was not as strong as the 2-fold stimulation of Ku. This suggests that in *P. aeruginosa*, the Ku C-terminal region is not essential for stimulating LigD polymerase activity, implying that the stimulation may be driven through Ku’s core domain or by an alternative interaction mechanism unique to this species.

### Critical LigD polymerase domain amino acids mediate Ku interaction, enzymatic activity and DNA binding

Alphafold 3 modelling suggested several amino acids that could mediate an interaction between Ku and the LigD polymerase domain. These included D162, V194, and R198, also identified by Morati et al., (2025) who found mutations to some of these amino acids critical for Ku-stimulated ligation (21), suggesting a potential role for these amino acids in the Ku–LigD interaction. To directly assess this, we used microscale thermophoresis (MST) and found that LigD mutants D162R, V194D, and R198E exhibited a 2- to 5-fold reduction in binding affinity for Ku. While the reduction in binding affinity was not statistically significant (p-values ranging from 0.17-0.3 compared to LigD), these residues have some contribution to the Ku-LigD complex formation **(Figure 8 AB, Supplemental Table 10, Supplemental Figure 10)**. In addition, we verified that all these LigD POL mutants had retained secondary structure similar to wildtype LigD via CD Spectroscopy **(Supplemental Figure 11)**.

**Figure 8:**
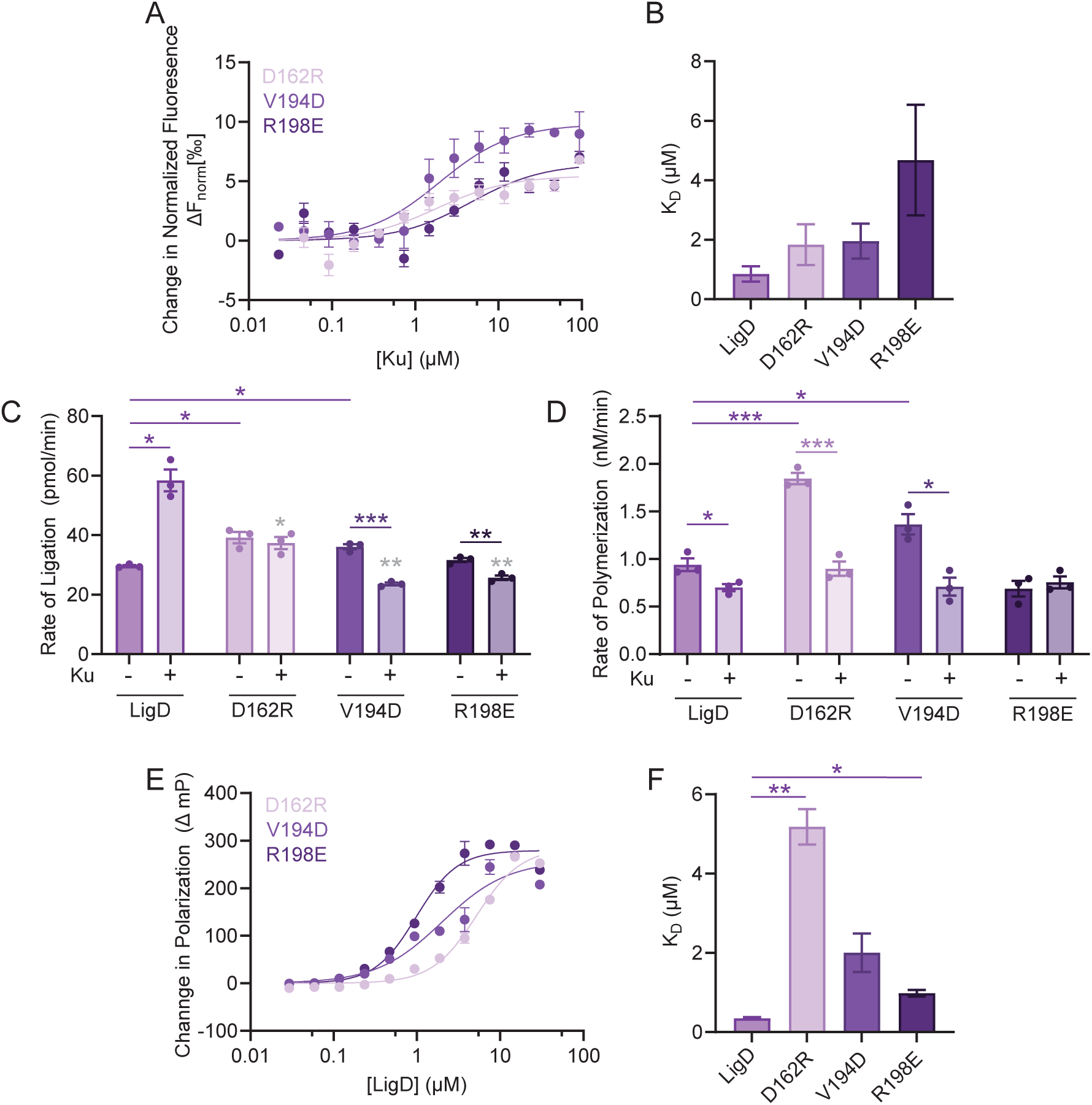
Key amino acids in the LigD polymerase domain modulate ligase activity. (A) Binding curves of the Ku-LigD interaction (LigD POL mutants) measuring the change in normalized fluorescence (ΔFnorm) of the LigD ligand over the concentration of Ku analyte (µM). Data is fitted with non-linear regression using the binding saturation – one site, specific function in Prism v.10.1.4 (GraphPad). (B) Apparent dissociation constants (KD) for Ku binding LigD and LigD POL mutants D162R, V194D and R198E. (C) Rates of Ligation of the linearized pUC19 plasmid by LigD and LigD mutants D162R, V194D and R198E in both the presence and absence of Ku. (D) Rates of polymerization of the 18-nucleotide overhang templated substrate by LigD and LigD mutants D162R, V194D and R198E in both the presence and absence of Ku. (E) DNA binding curves for LigD and LigD mutants D162R, V194D and R198E by fluorescence polarization. Data plotted is the change in polarization (mP) over the concentration of LigD (µM). (F) Apparent dissociation constants (KD) for LigD, D162R, V194D and R198E binding to the 40 bp DNA substrate. All data plotted are the mean ± standard error measure. n=3 technical replicates. ***p<0.001, **p<0.01, *p<0.05 (two-tailed t-test). Grey coloured * are in comparison to WT Ku+ WT LigD. MST raw traces are available in Supplemental Figure 10, ligation as a product of phosphate released is in Supplemental Figure 12 and representative gel images and curves of polymerization data are in Supplemental Figures 13 and 14.

We next quantitatively evaluated how these mutations affected LigD ligation activity. When tested in the absence of Ku, LigD D162R and V194D displayed slightly increased ligation rates compared to wild-type LigD, whereas R198E showed no significant change **(Figure 8 C, Supplemental Table 11, Supplemental Figure 12)**. However, in the presence of Ku, all three mutants exhibited ∼2-fold reduced ligation rates relative to wild-type Ku–LigD, aligning with findings by Morati et al. (21). Notably, for V194D and R198E, ligation activity in the presence of Ku dropped below levels observed with LigD alone, while D162R showed no change with or without Ku. These results support the conclusion that a reduced affinity for Ku impairs stimulation of LigD ligase activity, and confirms that D162, V194, and R198 are involved in this regulatory mechanism.

We also examined polymerase activity in the same mutants in the presence of Ku. For V194D and R198E, the loss of Ku interaction did not significantly alter polymerase activity relative to wild type Ku+LigD, reinforcing our earlier finding that Ku does not stimulate LigD template-dependent polymerase function **(Figure 8 D, Supplemental Table 12, Supplemental Figure 13-14)**. LigD D162R in the presence of Ku, also showed polymerase activity comparable to LigD alone, further supporting this observation. Interestingly, in the absence of Ku, both D162R and V194D mutants exhibited increased polymerase activity, with D162R showing a twofold enhancement relative to wild-type LigD. These findings indicate that specific POL domain mutations can enhance polymerase function, though further studies are needed to assess their DNA repair capacity *in vivo*.

To explore whether changes in enzymatic activity were linked to DNA binding, we measured the DNA binding affinity of each mutant **(Figure 8 EF, Supplemental Table 13)**. All three displayed a reduction in DNA binding compared to wild-type LigD. Notably, D162R and V194D, the mutants with elevated polymerase activity, had the most pronounced loss of DNA affinity (5.18±0.44 µM and 2.00±0.48 µM respectively), suggesting that these residues may contribute to DNA recognition and binding by the LigD polymerase domain.

### LigD ligase domain interactions contribute to the Ku-LigD binding interface

AlphaFold3 modelling of the Ku-LigD complex suggests that one monomer of Ku interacts with the polymerase domain of LigD through the Ku C-terminal tail, while the other Ku C-terminus interacts with the ligase domain of LigD, suggesting a Ku dimer could interact simultaneously with both the polymerase and ligase domains of LigD **(Figure 9, Supplemental Figure 15)**. Therefore, we wanted to investigate whether specific amino acid interactions between Ku and the LigD ligase domain also affect LigD enzymatic activity.

**Figure 9:**
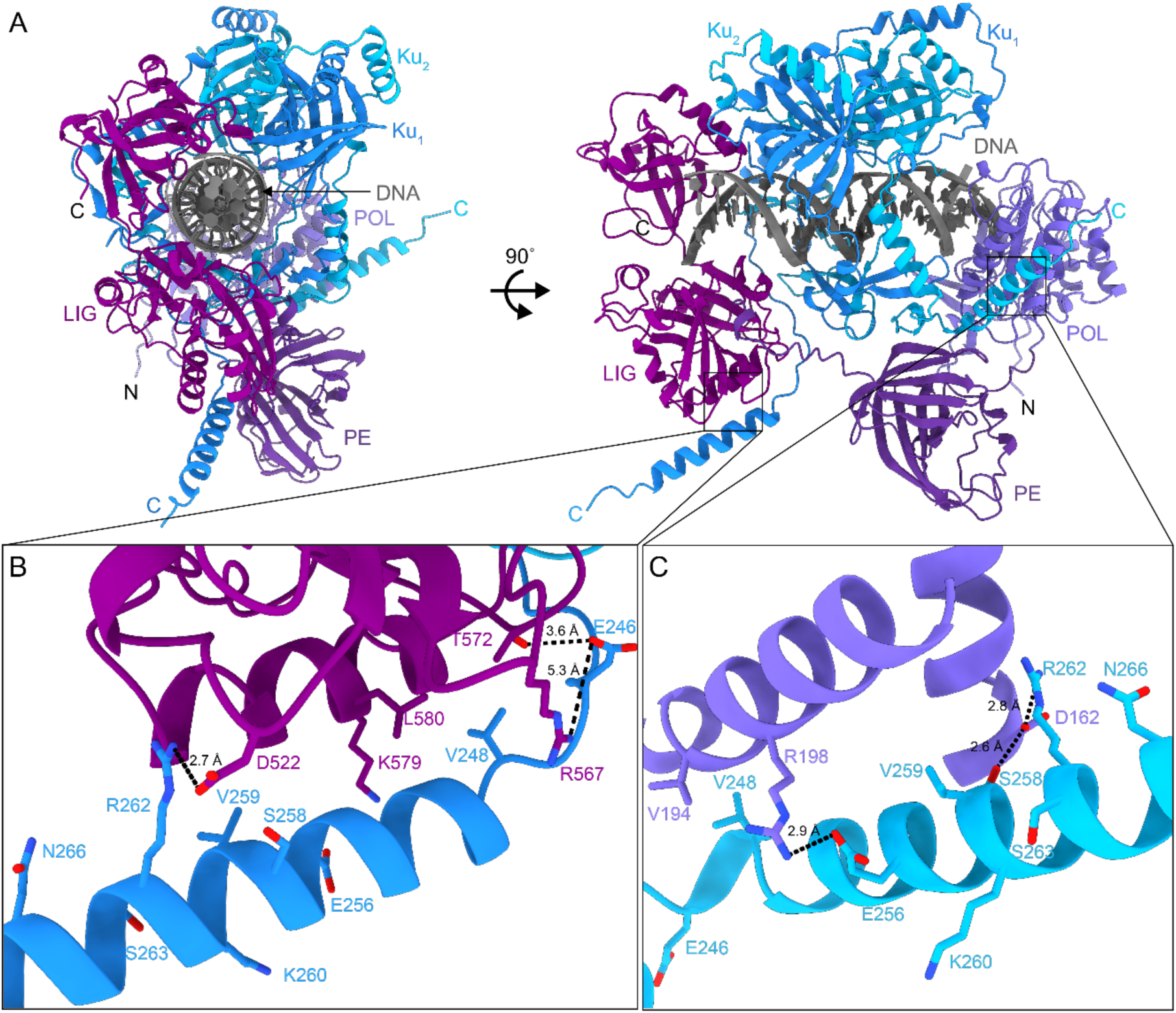
AlphaFold3 model of the Ku-LigD-DNA complex. (A) AlphaFold3 model of the Ku-LigD-DNA complex, showing DNA bound through central pore of Ku. Model was rotated 90° to show LigD binding the DNA and Ku. (B) Interactions between Ku C-terminus and LigD LIG domain. Potential hydrogen bond network between Ku and LigD are shown between Ku E246 and LigD T572/R567, with an additional hydrogen bond between Ku R262 and LigD D522. (C) Interactions between the Ku C-terminus and LigD POL domain. Potential hydrogen bond network between Ku and LigD are shown between Ku S258/R262 and LigD D162, with an additional hydrogen bond between Ku E256 and LigD R198. pLDDT and predicted error alignment (PAE) model are shown in Supplemental Figure 15.

We identified similar point mutations as Morati et al., (21) in the LigD ligase domain (D522R, K579E and L580E) that may play a role in the Ku-LigD interaction and verified that all mutants retained secondary structure similar to wildtype LigD using CD Spectroscopy **(Supplemental Figure 16)**. We then assessed the binding affinity of these LigD mutants to Ku through MST. We found that LigD D522R did not bind Ku strongly enough to calculate a KD under the same protein concentrations as the other LigD mutants, while LigD K579E had an increased binding affinity for Ku (KD = 0.51 ± 0.15 µM) and LigD L580E had decreased binding affinity for Ku (KD = 1.27 ± 0.15 µM) compared to wildtype LigD. **(Figure 10 AB, Supplemental Table 14, Supplemental Figure 10)**.

**Figure 10:**
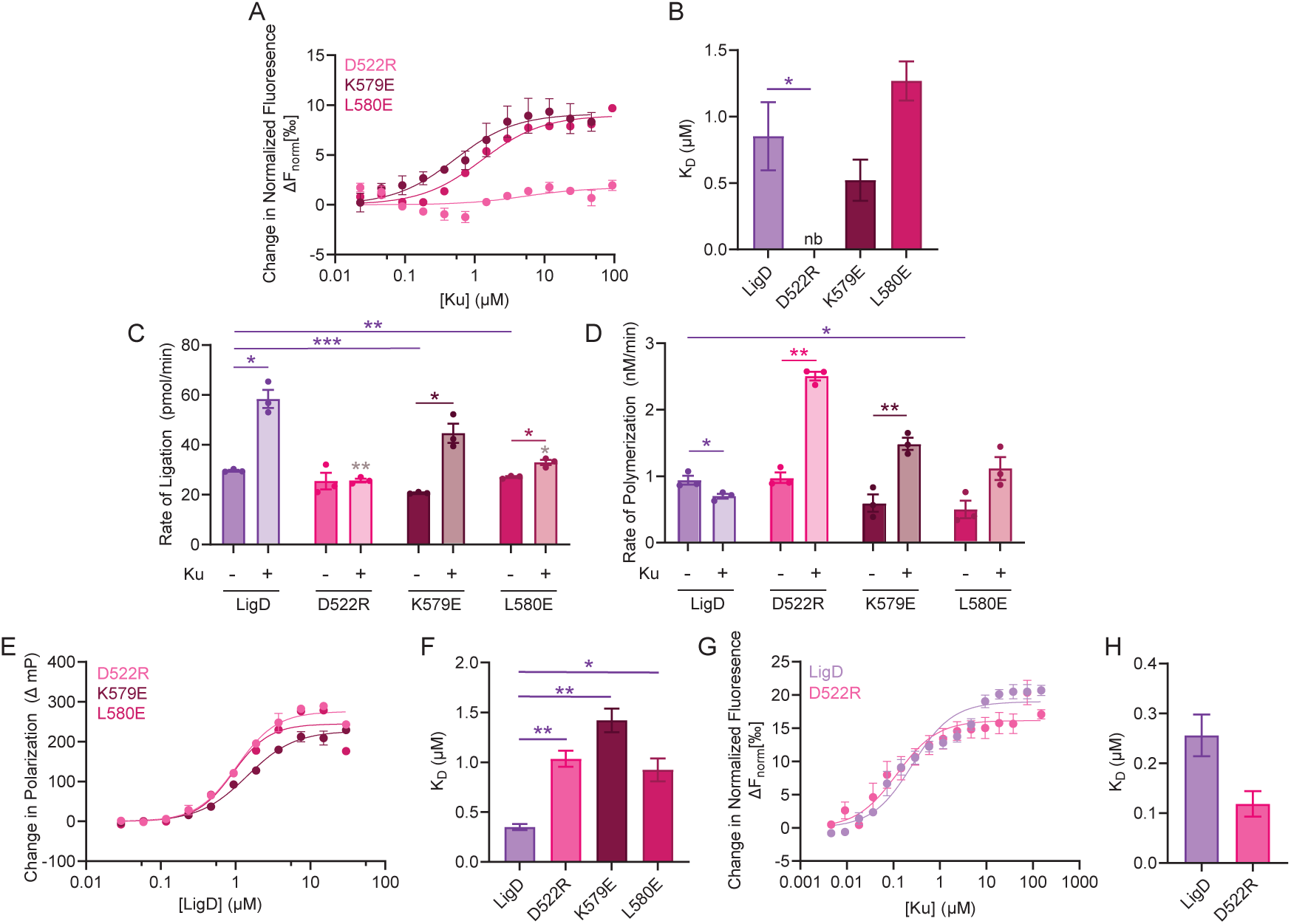
**Key amino acids in the ligase domain modulate LigD activity**. (A) Binding curves of the Ku-LigD interaction (LigD ligase mutants) measuring the change in normalized fluorescence (ΔFnorm) of the LigD ligand over the concentration of Ku analyte (µM). Data is fitted with non-linear regression using the binding saturation – one site, specific function in Prism v.10.1.4 (GraphPad). (B) Apparent dissociation constants (KD) for Ku binding LigD and LigD mutants D522R, K579E and L580E. nb – no binding (C) Rates of Ligation of the linearized pUC19 plasmid by LigD and LigD mutants D522R, K579E and L580E in both the presence and absence of Ku. (D) Rates of polymerization of the 18-nucleotide overhang templated substrate by LigD and LigD mutants D522R, K579E and L580E in both the presence and absence of Ku. (E) DNA binding curves for LigD and LigD mutants D522R, K579E and L580E by fluorescence polarization. Data plotted is the change in polarization (mP) over the concentration of LigD (µM). (F) Apparent dissociation constants (KD) for LigD, LigD D522R, K579E and L580E binding to the 40 bp DNA substrate. (G) Binding curves of the Ku-LigD-DNA ternary complexes with 40 bp dsDNA measuring the change in normalized fluorescence (ΔFnorm) of the LigD ligand over the concentration of DNA-bound Ku analyte (µM). Data is fitted with non-linear regression using the binding saturation – one site, specific function in Prism v.10.1.4 (GraphPad). (H) Apparent dissociation constants (KD) for the ternary complex formation between Ku-LigD-DNA and Ku-LigD D522R-DNA. All data plotted are the mean ± standard error measure. n=3 technical replicates. ***p<0.001, **p<0.01, *p<0.05 (two-tailed t-test) in comparison to LigD (purple lines) or LigD mutants (pink lines). Light brown coloured * are in comparison to WT Ku+ WT LigD. MST raw traces are available in Supplemental Figure 10, ligation as a product of phosphate released is in Supplemental Figure 17 and representative gel images and curves of polymerization data are in Supplemental Figures 18-19. MST raw traces are available in Supplemental Figure 20 for ternary complexes in Figure 10 GH.

We then wanted to investigate the ligation activity of these LigD mutants further using our ligation assay to compare rates of ligation with the ligation activities reported (21). The ligation rates of LigD mutants D522R, K579E and L580E in the absence of Ku were similar to wild-type LigD **(Figure 10 C, Supplemental Table 15, Supplemental Figure 17)**. Upon addition of Ku, LigD D522R showed no rate change as expected, given it did not bind Ku. However, LigD K579E had a 2-fold increase in ligation rate and LigD L580E showed a slight increase in ligation rate compared to wildtype LigD activity in the absence of Ku, although these levels were decreased compared to ligation by wildtype LigD with Ku. These findings differ slightly from Morati *et al*., for LigD D522R and K579E, where they observed increased ligation (21). However, differences in experimental parameters (1kb blunt-ended vs. 2.7kb sticky-ended ligation substrate; protein concentrations; methodology) may account for observed differences.

To better understand if there may be a potential allosteric inhibition on polymerase activity mediated by interactions between Ku and the LigD ligase domain, we examined the polymerase activity of the LigD ligase domain mutants. We found that LigD D522R had a polymerization rate comparable to wild-type LigD, while LigD K579E and LigD L580E had 1.5-fold and 7-fold decreased rates of polymerization, respectively, in comparison to wild-type LigD **(Figure 10D, Supplemental Table 16, Supplemental Figures 18-19)**. Surprisingly, the addition of Ku stimulated polymerase activity in these mutants 1.5 – 3-fold higher than that of Ku and LigD, and of LigD alone. To determine whether an increased DNA binding affinity would explain the Ku-stimulated polymerase activity, we measured the binding affinity between a 40 bp DNA substrate and the LigD ligase domain mutants **(Figure 10 EF, Supplemental Table 17)**. We found that the LigD ligase domain mutants all had a 3-fold decrease in DNA binding affinity compared to wild-type LigD, which would not correlate with an increase in polymerase activity. Therefore, we checked the affinity of the ternary complex and whether the presence of Ku, DNA and the LigD ligase mutants resulted in a strong binding interaction. Interestingly, the affinity between a DNA-bound Ku and LigD D522R was 0.12 ± 0.025 µM **(Figure 10 GH, Supplemental Table 18, Supplemental Figure 20)**, indicative of a strong interaction. It is possible that LigD D522R combined with Ku arranges the DNA in a highly favourable conformation for polymerase activity compared to wildtype LigD and Ku, although further experiments are needed to confirm this hypothesis.

## Discussion

Previous studies have established that Ku enhances the enzymatic activities of LigD, with *M. tuberculosis* Ku stimulating ligation and *P. aeruginosa* Ku stimulating both ligation and polymerization (6, 22); however, the molecular details of this interface remain undefined. In this study, we applied a structure-informed approach to characterize the Ku-LigD interaction, providing mechanistic insights into Ku-mediated repair activities associated with NHEJ. Here, we find that the Ku C-terminal region makes important contacts with both the polymerase and ligase domains of LigD that correlate with enhanced ligation and limited polymerization.

Due to the absence of an experimental structure of the Ku-LigD-DNA complex, we used AlphaFold3 modelling (23) to predict the architecture of the *M. tuberculosis* Ku-LigD complex bound to DNA **(Figure 11)**. Consistent with the reported cryo-EM structures of Ku bound to DNA, the core domain of the Ku homodimer threads onto DNA, forming a tightly bound complex to DNA ends (24–26). Notably, this AlphaFold3 model predicts that the C-terminus of one Ku monomer contacts the polymerase domain of LigD, while the second C-terminus engages the ligase domain, suggesting a bipartite interaction interface **(Figure 9 BC)**. This model aligns with our previous biochemical data showing that a Ku homodimer could interact with both the LigD ligase and LigD polymerase domains(6).

**Figure 11:**
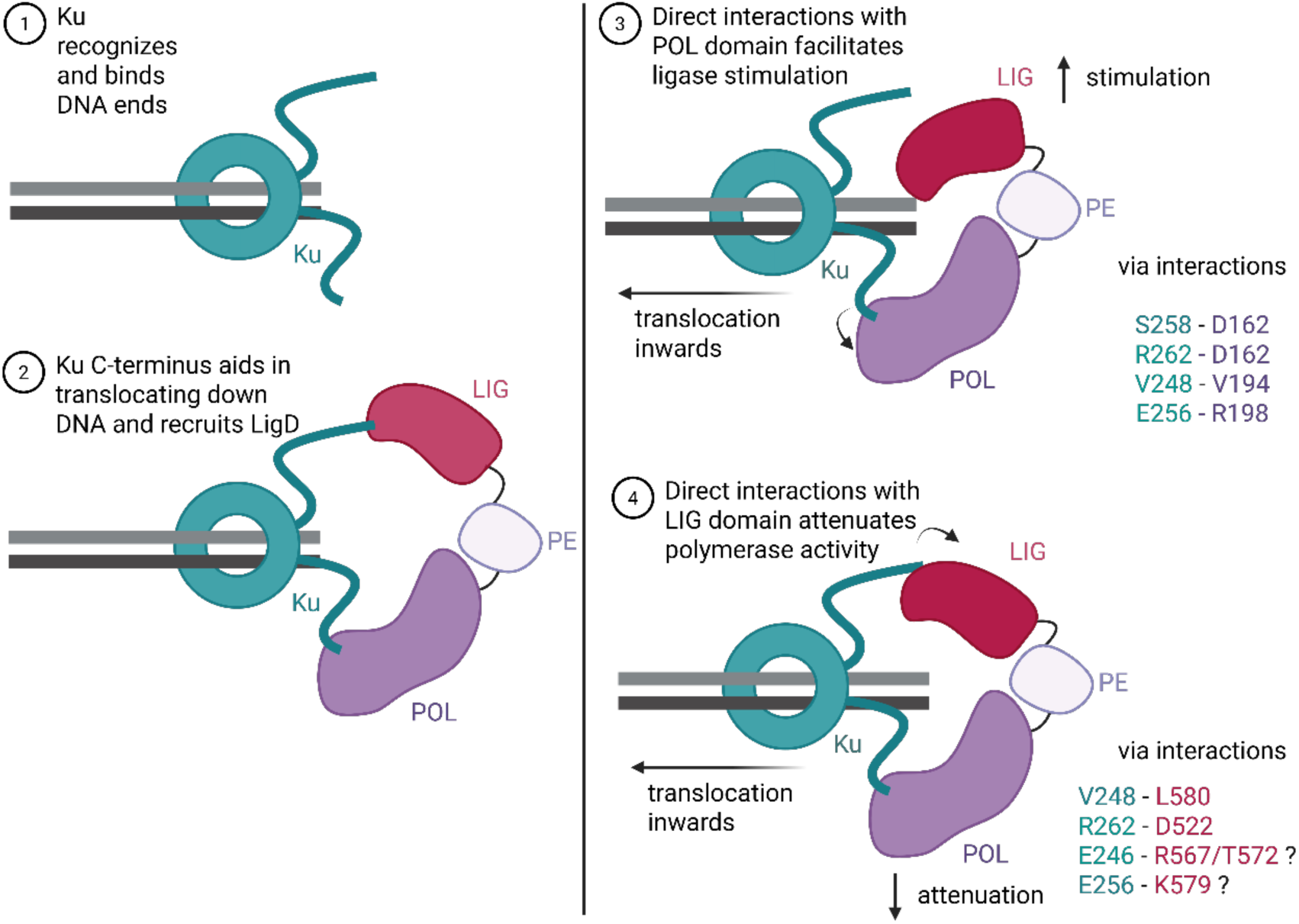
Proposed model for Ku C-terminal regulation of LigD activity during mycobacterial NHEJ. (1) Ku recognizes and binds the DNA ends of the break via the core region, with the flexible C-terminus available to recruit LigD (only one side of the DNA break shown for clarity). (2) Ku C-terminus recruits LigD to the break. LigD is composed of a POL (purple), PE (light purple), and LIG (pink) domains. Ku C-termini interact with both the POL and LIG domains to facilitate LigD activity. (3) Direct interactions between the Ku C-terminus and the POL domain (Ku S258 – LigD D162, Ku R262 – LigD D162 and Ku E256 – LigD R198) facilitate LigD ligase stimulation. (4) Direct interactions between the Ku C-terminus and the LIG domain (Ku R262 – LigD D522, with potential interactions between Ku E246 – LigD R567/T572, and Ku E256 – LigD K579) facilitates LigD polymerase attenuation.

Consistent with this structural model, our combined NMR and MST protein-binding studies **(Figure 2)** identified five key residues in the Ku C-terminus (E246, V248, S258, K260 and N266) predicted to interact with the LigD polymerase domain. Functionally, we found these amino acids, along with Ku 258 previously (6), are needed for high rates of ligation **(Figure 3)**, as mutations to these amino acids abolish or reduce Ku-stimulated ligation. Interestingly, AlphaFold3 suggests that of these residues, Ku V248 and S258 form clear interactions with the polymerase domain, while Ku E246 and S258 could interact with the ligase domain, dependent on movement in the Ku C-terminus to bring it closer to LigD **(Figure 9 BC)**. LigD residues in both the polymerase (D162, V194, R198) and the ligase (D522, K579, L580) domains reported to be involved in Ku binding (21), and quantified here **(Figure 9)**, are also seen in the model, aligning with separate Ku C-terminal tails. Between the Ku C-terminus and the LigD polymerase domain, the AlphaFold3 model suggests that LigD D162 forms a hydrogen bond network with Ku S258 and R262, with additional hydrogen bonding between LigD R198 and Ku E256. LigD V194 and Ku V248 contribute to Van der Waals packing at the helix-helix interface. In the ligase domain, the AlphaFold3 model suggests potential hydrogen-bond interactions between LigD D522-Ku R262, while LigD L580 contributes to the hydrophobic interface of a helical bundle with Ku V248. These contacts all support a bipartite interaction mechanism between Ku and LigD. The AlphaFold3 model mostly aligns with our protein-protein interaction studies **(Figures 2, 9)**, although depending on flexibility and movement of the Ku C-terminus, it can be seen how the other residues identified (ie. Ku E256, K260, N266) may also interact with LigD. These findings collectively demonstrate that direct interactions between the Ku C-terminus and LigD are essential for Ku-mediated ligase stimulation, aligning with results by Morati et al., (21) and refining the molecular model of how Ku regulates LigD function.

While prior studies in *P. aeruginosa* have shown that Ku enhances LigD template-dependent polymerase activity (12), our findings revealed that this mechanism is not conserved in *M. tuberculosis*. Instead, we observed that the addition of Ku led to a modest, but consistent attenuation of LigD template-dependent polymerase activity, with this attenuation being more pronounced when Ku lacks the C-terminal region (Kucore) **(Figures 4-5, 7)**. An attenuated polymerase activity may be a mechanism for *M. tuberculosis* to limit mutations introduced by the error-prone polymerase domain, as fewer mutations would be beneficial for a slower growing bacteria in a stable environment and may explain why the Ku C-terminal region of *M. tuberculosis* and *P. aeruginosa* differ greatly in their sequence composition (27, 28). Of the five amino acids identified in this study to interact with LigD, only S258 is strongly conserved, while V248 and K260 (I248 and R260 in *P. aeruginosa* respectively) conserve similar amino acid side chains (6, 7). From unpublished work in our lab, we have also measured the interaction affinity between *P. aeruginosa* Ku and LigD and found that it is weaker than the interaction of the *M. tuberculosis* homologs, which may be attributed to these differences at key residues in the Ku C-terminus. Overall, these findings suggest a species-specific regulatory divergence in NHEJ.

This attenuation of polymerase activity appears to be mediated by specific interactions between the Ku C-terminus and LigD, as disrupting residues E246, S258, and N266 restored polymerase activity levels comparable to LigD alone **(Figure 5)**. These results further support a model where direct *M. tuberculosis* Ku-LigD contacts modulate polymerase activity, in contrast to the stimulatory role observed in *P. aeruginosa* homologs. Interestingly, Ku V248 did not influence polymerase activity, indicating that not all residues that are critical for the Ku-LigD interaction play a role in polymerase regulation. Unexpectedly, Ku K260A led to a significant decrease in polymerase activity, similar to the Kucore protein **(Figures 5, 7)**, contradicting our hypothesis that a loss of interaction would restore LigD activity. Previous work has shown that the Ku C-terminal tail is needed to translocate along DNA (6, 14, 22) . We also know that there is a high affinity between DNA and Kucore (6), while studies from our lab using atomic force microscopy show Kucore trapped at DNA ends, and similar to the results here, prevent Ku-stimulated UvrD1 helicase activity (Warner et al., manuscript in revision), illustrating how without the C-terminal tail, Kucore is unable to translocate, which is likely the cause of the significantly reduced polymerase activity observed in Figure 7. We propose that a similar effect is occurring with Ku K260A, where Ku K260 may be necessary for Ku translocation along DNA, and when mutated, Ku remains bound to DNA ends. Our DNA binding studies further support this hypothesis with Ku K260A having an 8-fold increase in affinity for a dsDNA substrate compared to wild-type Ku, although how Ku K260A interacts with DNA based on the AlphaFold3 model is unclear **(Figure 6, 9)**.

Previously, we and others have suggested that interactions between Ku and the LigD polymerase domain were necessary to stimulate ligation activity, but we had also seen direct interactions between Ku and the ligase domain (4, 6, 13, 14, 22, 29). We dissected this mechanism of Ku-stimulated ligation further by evaluating amino acids in the polymerase domain critical to this interaction and activity **(Figure 8)** but also discovered here that interactions between Ku and the LigD ligase domain are necessary for Ku-stimulated ligation **(Figure 10)**. In particular, LigD D522R and L580E had very weak interactions with Ku and severely reduced Ku-stimulated ligation. These findings suggest that while the polymerase domain may serve as the primary interface between Ku and LigD to enhance ligation, specific residues within the ligase domain also contribute to this regulatory mechanism, stabilizing the Ku-LigD complex.

In summary, our findings define a dual regulatory role for the Ku C-terminal region in modulating LigD enzymatic activity during NHEJ **(Figure 11)**. We propose that in NHEJ, Ku recognizes and binds the DNA ends, while the Ku C-terminus allows Ku to translocate down the DNA to make room for LigD at the DNA break. Ku then recruits and stimulates LigD ligase activity through specific interactions with both the polymerase and ligase domains, while concurrently attenuating polymerase activity, likely via modulating the conformation of the proteins on DNA. These observations contrast with the purely stimulatory activity observed in *P. aeruginosa* homologs for both polymerase and ligase activity, highlighting a mechanistic divergence in bacterial NHEJ regulation. Furthermore, our results highlight the complexity of Ku’s role as both a structural scaffold and an active regulator of LigD enzymatic functions, mediated through a combination of protein-protein and protein-DNA interactions. The current structures of bacterial Ku homologs bound to DNA offer the first insights into important Ku-DNA interactions, but are missing the C-terminus, likely due to its flexibility, which is needed for function (24–26). Future structural studies of a Ku-LigD-DNA complex will be informative to this model. Together, these insights enhance our understanding of the molecular basis of Ku-LigD cooperation in NHEJ, illustrating how species-specific adaptations fine-tune DNA repair mechanisms, and may identify potential antimicrobial targets aimed at inhibition of the NHEJ repair machinery.

## Acknowledgements

Results shown in this report were obtained from work performed in the Centre for Microbial Chemical Biology, at McMaster University. We acknowledge the help of Dr. Giuseppe Melacini and Dr. Madoka Akimoto. D.J.S., was supported by a National Sciences and Engineering Research Council of Canada with a Canadian Graduate Scholarship (CGS-D). We thank Dr. Fredrik Westerlund and Dr. Florian Morati at Chalmers University of Technology for thoughtful discussions of the results. BioRender was used to create the following Images: Graphical Abstract (Created in BioRender. Sowa, D. (2025) https://BioRender.com/c5tk28h) and Final Model (Created in BioRender. Sowa, D. (2025) https://BioRender.com/ueey2nw). Protein models were generated using ChimeraX (version 1.9) (30).

