## Supplemental Data for "The *Mycobacterium tuberculosis* Ku C-terminus orchestrates LigD activity through domain-specific interactions"

**Supplementary Material**

**Supplemental Table 1:** Cloning Primers for Protein Expression Plasmids

| Protein Expression Construct | Oligonucleotides | Plasmid Backbone | Cloning Method |
|---|---|---|---|
| Ku E246A | 5' TGACGAGCCGGCGGATGTGTCGG 3'<br>5' AGTAAGCGAGGCTGATCCTCG 3' | pMCSG7 | Q5-SDM |
| Ku V248A | 5' GCCGGAGGATGCGTCGGATTTAC 3'<br>5' TCGTCAAGTAAGCGAGGC 3' | pMCSG7 | Q5-SDM |
| Ku E256A | 5'GTTAGCAGCGAGTGTCAAAGCCCGTTCCAAAGCTAACTCTAATGTTCCGACCCC 3'<br>5' CTCGCTGCTAACTTTGCAAGTAAATCCGACACATCCTCCGGCTCGTCA AGTAAGCG 3' | pMCSG7 | SDM |
| Ku V259A | 5' GCGAGTGCGAAAGCCCGTTCCAAAGCTAACTCTAATGTTCCG 3'<br>5' GGCTTTCGCACTCGCTTCTAACTTTGCAAGTAAATCCGACAC 3' | pMCSG7 | SDM |
| Ku K260A | 5' AGCGAGTGTCGCGGCCCGTTCAAAG 3'<br>5' TCTAACTTTGCAAGTAAATCCG 3' | pMCSG7 | Q5-SDM |
| Ku A261E | 5' GAGTGTCAAAGAGCGTTCCAAAG 3'<br>5' GCTTCTAACTTTGCAAGTAAATC 3' | pMCSG7 | Q5-SDM |
| Ku S263A | 5' CAAAGCCCGTGCCAAAGCTAACT 3'<br>5' ACACTCGCTTCTAACTTTGC 3' | pMCSG7 | Q5-SDM |
| Ku N266A | 5' TTCCAAAGCTGCCTCTAATGTTCCGACCC 3'<br>5' CGGGCTTTGACACTCGCT 3' | pMCSG7 | Q5-SDM |
| LigD D162R | 5' CTTGCTCGCATTGGCCTGGTTACCTTTCCGGTGACATCGGGCTCAAAAG 3'<br>5' CCAATGCGAGCAAGCAGGTCACGAACAGCGCGTGCTACCTCAG 3' | pMCSG7 | SDM |
| LigD V194D | 5'CTACTGATCTTGCTAAGCGTGTGGCTCAACGTTTAGAACAGGCAATGCCGGCTTTG 3'<br>5' GCAAGATCAGTAGCTCCACGGGATGAGACAGGTTCATCCAGGGGAG 3' | pMCSG7 | SDM |
| LigD R198E | 5' CTAAGGAAGTGGCTCAACGTTTAGAACAGGCAATGCCGGCTTTGGTGAC 3'<br>5' GCCACTTCCTTAGCAAGAACAGTAGCTCCACGGGATGAGACAGG 3' | pMCSG7 | SDM |
| LigD D522R | 5' CTGGCTCGTCATCATGTCGTACTGGATGGAGAGGCAGTAGTTCTGGACAGTAG 3'<br>5' GATGACGAGCCAGGTCTTCTGCAAGAGCACGCAATTGCGGGTATTCAGCTG 3' | pMCSG7 | SDM |
| LigD K579E | 5' CGTCGCGAGCTGTTGGAGACTCTGGCGAACGCAACTTCATTGACGG 3'<br>5' CTCCAACAGCTCGCGACGATCCTGATAACGAGTTCCTAAAAGTGCGCGACC 3' | pMCSG7 | SDM |
| LigD L580E | 5' CGTCGCAAGGAGTTGGAGACTCTGGCGAACGCAACTTCATTGACGG 3'<br>5' CTCCAACTCCTTGCGACGATCCTGATAACGAGTTCCTAAAAGTGCGCGACC 3' | pMCSG7 | SDM |

**Supplemental Table 2:** Oligonucleotide sequences for DNA binding and enzyme assay substrates.

| Figure Panel | Substrate | Oligonucleotides |
|---|---|---|
| **6 A-B, 9 E-F, 11 E-F** | 40bp dsDNA | 5' 6-FAM CAGGGTAAGTGTGGAGGTGTAGGGAAGGGAATGTTGTCTG 3'<br>5'CAGACAACATTCCCTTCCCTACACCTCCACACTTACCCTG 3' |
| **4 A-E, 5 A-B, 6 C-D, 9 D, 11 D** | 18nt overhang DNA | 5' (PO4) CAACTGCAGTTCTAGACC 6-FAM3'<br>5' GATGTAATCCCTCGATGAGGTCTAGAACTGCAGTTG 3' |

**Supplemental Table 3:** Molecular Weight and Oligomeric State of Ku mutants by SEC-MALS (data from supplemental figure 3).

| Protein | Theoretical Molecular Weight (kDa) | Multi-Angle Light Scattering Molecular Weight (kDa) | Oligomeric State |
|---|---|---|---|
| Ku E246A | 33.7 | 65.3 | Dimer |
| Ku V248A | 33.7 | 64.7 | Dimer |
| Ku E256A | 33.7 | 63.1 | Dimer |
| Ku V259A | 33.7 | 69.2 | Dimer |
| Ku K260A | 33.7 | 71.3 | Dimer |
| Ku A261E | 33.7 | 60.4 | Dimer |
| Ku S263A | 33.7 | 65.5 | Dimer |
| Ku N266A | 33.7 | 76.9 | Dimer |

**Supplemental Table 4:** Ku point mutant-LigD Binding Affinity $K_D$ Values (data from figure 2). p-value, compared to Ku-LigD interaction.

| Ligand | Analyte | $K_D$ ± SEM (μM) | p-value |
|---|---|---|---|
| Ku | LigD | 0.85 (± 0.26) | - |
| Ku E246A | LigD | 10.07 (± 2.88) | 0.0841 |
| Ku V248A | LigD | 4.22 (± 1.63) | 0.1718 |
| Ku E256A | LigD | 0.28 (± 0.03) | 0.1561 |
| Ku S258A | LigD | 17.22 (± 4.50) | 0.0673 |
| Ku V259A | LigD | 0.79 (± 0.14) | 0.8487 |
| Ku K260A | LigD | 6.25 (± 0.91) | 0.0210 |
| Ku A261E | LigD | 0.87 (± 0.22) | 0.9526 |
| Ku R262A | LigD | 0.26 (± 0.02) | 0.1464 |
| Ku S263A | LigD | 0.35 (± 0.10) | 0.1812 |
| Ku N266A | LigD | no-binding (nb) | 0.0801 |

**Supplemental Table 5:** Ku-stimulated ligation rates (data from figure 3). p-value, protein indicated in parentheses indicates comparator.

| Protein 1 | Protein 2 | Rate ± SEM (pmol/min) | p-value (LigD) | p-value (Ku+LigD) |
|---|---|---|---|---|
| LigD | Ku E246A | 22.84 (± 1.30) | 0.0280 | 0.0054 |
| LigD | Ku V248A | 35.63 (± 5.63) | 0.3975 | 0.0350 |
| LigD | Ku K260A | 43.62 (± 1.03) | 0.0025 | 0.0470 |
| LigD | Ku N266A | 30.45 (± 1.80) | 0.6906 | 0.0068 |

**Supplemental Table 6:** Templated addition polymerization rates for LigD (data from figure 4). p-value, protein indicated in parentheses indicates comparator.

| Protein 1 | Protein 2 | Rate ± SEM (nM/min) | p-value (LigD) |
|---|---|---|---|
| LigD | - | 0.94 (±0.07) | - |
| LigD | Ku | 0.70 (± 0.04) | 0.0493 |

**Supplemental Table 7:** Templated addition polymerization rates for LigD and Ku mutants (data from figure 5). p-value, protein indicated in parentheses indicates comparator.

| Protein 1 | Protein 2 | Rate ± SEM (nM/min) | p-value (LigD) | p-value (Ku) | p-value ($Ku_{core}$) |
|---|---|---|---|---|---|
| LigD | $Ku_{core}$ | 0.27 (± 0.01) | 0.0091 | 0.0055 | - |
| LigD | Ku E246A | 1.03 (± 0.05) | 0.3629 | 0.0063 | - |
| LigD | Ku V248A | 0.65 (± 0.11) | 0.0933 | 0.6788 | - |
| LigD | Ku S258A | 0.90 (± 0.09) | 0.7237 | 0.1543 | - |
| LigD | Ku K260A | 0.26 (± 0.03) | 0.0045 | 0.0011 | 0.7074 |
| LigD | Ku N266A | 0.85 (± 0.08) | 0.4620 | 0.1827 | - |

**Supplemental Table 8:** Ku-DNA binding affinity apparent $K_D$ values (data from figure 6). p-value, protein indicated in parentheses indicates comparator. Wildtype values are from (6).

| Protein | 40bp | p-value (Ku) |
|---|---|---|
| Ku E246A | 0.46 (± 0.08) | 0.0645 |
| Ku V248A | 3.19 (± 0.46) | 0.9617 |
| Ku K260A | 0.37 (± 0.03) | 0.0640 |
| Ku N266A | 3.62 (± 0.52) | 0.6314 |

**Supplemental Table 9:** Templated addition polymerization rates of *P. aeruginosa* LigD (data from figure 7). p-value, protein indicated in parentheses indicates comparator.

| Protein 1 | Protein 2 | Rate ± SEM (nM/min) | p-value (LigD) | p-value (Ku) |
|---|---|---|---|---|
| LigD | - | 0.65 (±0.02) | - | - |
| LigD | Ku | 1.29 (± 0.05) | 0.0031 | - |
| LigD | $Ku_{core}$ | 0.89 (±0.02) | 0.0019 | 0.0066 |

**Supplemental Table 10:** Ku-LigD polymerase domain point mutants binding affinity apparent $K_D$ values (data from figure 8A-B). p-value, protein indicated in parentheses indicates comparator.

| Ligand | Analyte | $K_D$ ± SEM (μM) | p-value (LigD) |
|---|---|---|---|
| LigD D162R | Ku | 1.84 (± 0.69) | 0.2869 |
| LigD V194D | Ku | 1.95 (± 0.59) | 0.1946 |
| LigD R198E | Ku | 4.68 (± 1.86) | 0.1734 |

**Supplemental Table 11:** Ligation rates for LigD polymerase domain point mutants. p-value protein indicated in parentheses indicates comparator (data from figure 8C). Wildtype Ku values from (6).

| Protein 1 | Rate ± SEM (pmol/min) | p-value (LigD) | + Protein 2 | Rate ± SEM (pmol/min) | p-value (+Ku) | p-value (WT Ku+ WT LigD) |
|---|---|---|---|---|---|---|
| LigD D162R | 39.18 (± 1.89) | 0.0332 | Ku | 36.34 (± 2.03) | 0.5440 | 0.0135 |
| LigD V194D | 36.08 (± 0.85) | 0.0086 | Ku | 23.58 (± 0.48) | 0.0008 | 0.0098 |
| LigD R198E | 31.65 (± 0.66) | 0.0708 | Ku | 25.69 (± 0.74) | 0.0039 | 0.0099 |

**Supplemental Table 12:** Templated addition polymerization rates of LigD polymerase domain point mutants (data from figure 8D). p-value, protein indicated in parentheses indicates comparator. Wild-type Ku+LigD polymerase rate from supplemental table 6.

| Protein 1 | Rate ± SEM (nM/min) | p-value (LigD) | + Protein 2 | Rate ± SEM (nM/min) | p-value (Ku+LigD) |
|---|---|---|---|---|---|
| LigD D162R | 1.85 (± 0.07) | 0.0006 | Ku | 0.90 (± 0.09) | 0.0007 |
| LigD V194D | 1.36 (± 0.11) | 0.0368 | Ku | 0.71 (± 0.09) | 0.0104 |
| LigD R198E | 0.69 (± 0.08) | 0.0797 | Ku | 0.75 (± 0.06) | 0.5574 |

**Supplemental Table 13:** LigD polymerase domain point mutants DNA binding apparent $K_D$ values (data from figure 8E-F). p-value, protein indicated in parentheses indicates comparator.

| Protein | $K_D$ ± SEM (μM) | p-value (LigD) |
|---|---|---|
| LigD | 0.35 (± 0.03) | - |
| LigD D162R | 5.18 (± 0.44) | 0.0083 |
| LigD V194D | 2.00 (± 0.48) | 0.3832 |
| LigD R198E | 0.98 (± 0.08) | 0.0696 |

**Supplemental Table 14:** Ku-LigD ligase domain point mutants binding affinity $K_D$ values (data from figure 10A-B). p-value, protein indicated in parentheses indicates comparator. Wild-type Ku+LigD interaction from supplemental table 4.

| Ligand | Analyte | $K_D$ ± SEM (μM) | p-value (LigD) |
|---|---|---|---|
| LigD D522R | Ku | no-binding (nb) | 0.0801 |
| LigD K579E | Ku | 0.51 (± 0.15) | 0.3448 |
| LigD L580E | Ku | 1.27 (± 0.15) | 0.2484 |

**Supplemental Table 15:** Ligation Rates for LigD ligase domain point mutants (data from figure 10C). p-value, protein indicated in parentheses indicates comparator.

| Protein 1 | Rate ± SEM (pmol/min) | p-value (LigD) | + Protein 2 | Rate ± SEM (pmol/min) | p-value (+Ku) | p-value (WT Ku+ WT LigD) |
|---|---|---|---|---|---|---|
| LigD D522R | 25.50 (± 3.35) | 0.3423 | Ku | 25.70 (± 0.74) | 0.9560 | 0.0099 |
| LigD K579E | 20.70 (± 0.74) | 0.0002 | Ku | 44.60 (± 3.88) | 0.0250 | 0.0614 |
| LigD L580E | 27.10 (± 0.35) | 0.0071 | Ku | 32.90 (± 1.06) | 0.0228 | 0.0141 |

**Supplemental Table 16:** Templated addition polymerization rates of LigD ligase domain point mutants (data from figure 10D). p-value, protein indicated in parentheses indicates comparator.

| Protein 1 | Rate ± SEM (nM/min) | p-value (LigD) | + Protein 2 | Rate ± SEM (nM/min) | p-value (+Ku) |
|---|---|---|---|---|---|
| LigD D522R | 0.98 (± 0.08) | 0.7349 | Ku | 2.58 (± 0.15) | 0.0002 |
| LigD K579E | 0.60 (± 0.13) | 0.0987 | Ku | 1.49 (± 0.09) | 0.0067 |
| LigD L580E | 0.45 (± 0.10) | 0.0139 | Ku | 1.10 (± 0.17) | 0.0438 |

**Supplemental Table 17:** LigD ligase domain point mutants DNA binding apparent $K_D$ values (data from figure 10E-F). p-value, protein indicated in parentheses indicates comparator.

| Protein | $K_D$ ± SEM (μM) | p-value (LigD) |
|---|---|---|
| LigD | 0.35 (± 0.03) | - |
| LigD D522R | 1.03 (± 0.08) | 0.0915 |
| LigD K579E | 1.42 (± 0.12) | 0.8436 |
| LigD L580E | 0.92 (± 0.12) | 0.0540 |

**Supplemental Table 18:** LigD ligase domain point mutant D522R ternary complex formation (data from figure 10G-H). p-value, protein indicated in parentheses indicates comparator.

| Ligand | Analyte | $K_D$ ± SEM (μM) | p-value (LigD) |
|---|---|---|---|
| LigD | Ku | 0.25 (± 0.04) | - |
| LigD D522R | Ku | 0.12 (± 0.03) | 0.3369 |

**Supplemental Figures**

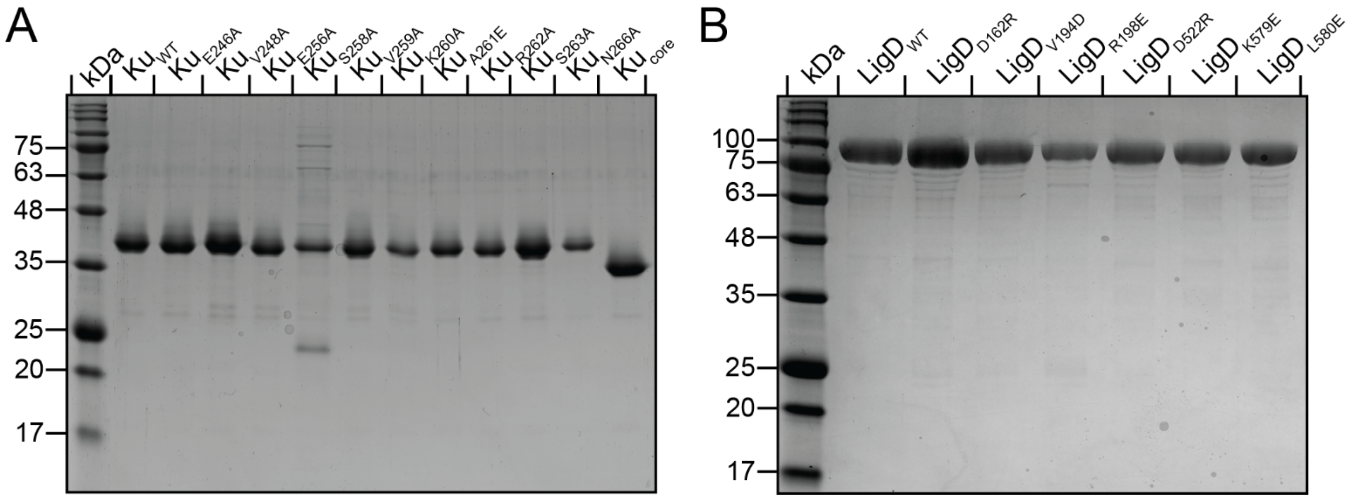


**Supplemental Figure 1:** Purified proteins used throughout this study. (A) Purified Ku, $Ku_{core}$ and Ku mutant proteins. (B) Purified LigD and LigD mutant proteins. 2 μg of each purified protein was loaded onto a 12% SDS-PAGE gel, run at 180V for 60 minutes.

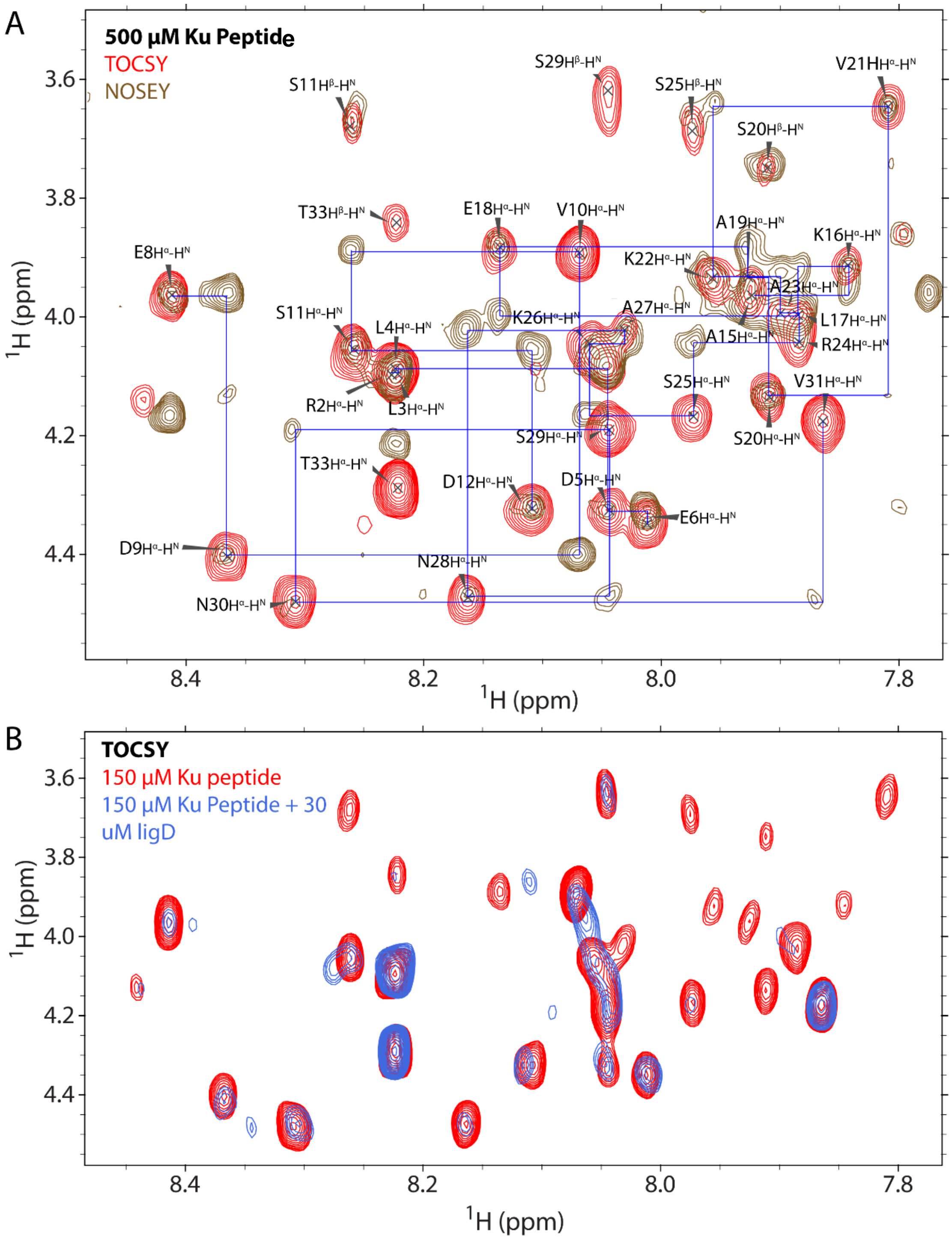


**Supplemental Figure 2.** Zoomed-in region of 2D spectra of the Ku peptide. (A) Overlay of $^{1}H$-$^{1}H$ TOCSY (red) and $^{1}H$-$^{1}H$ NOESY (brown). The assigned residues are labeled with the amino acid and corresponding atoms. The blue lines indicate the sequential connection between adjacent residues. Ambiguous assignments for the residues are removed. (B) Overlay of $^{1}H$-$^{1}H$ TOCSY spectra of the Ku peptide in the absence of LigD polymerase domain (red) and in the presence of LigD polymerase domain (blue).

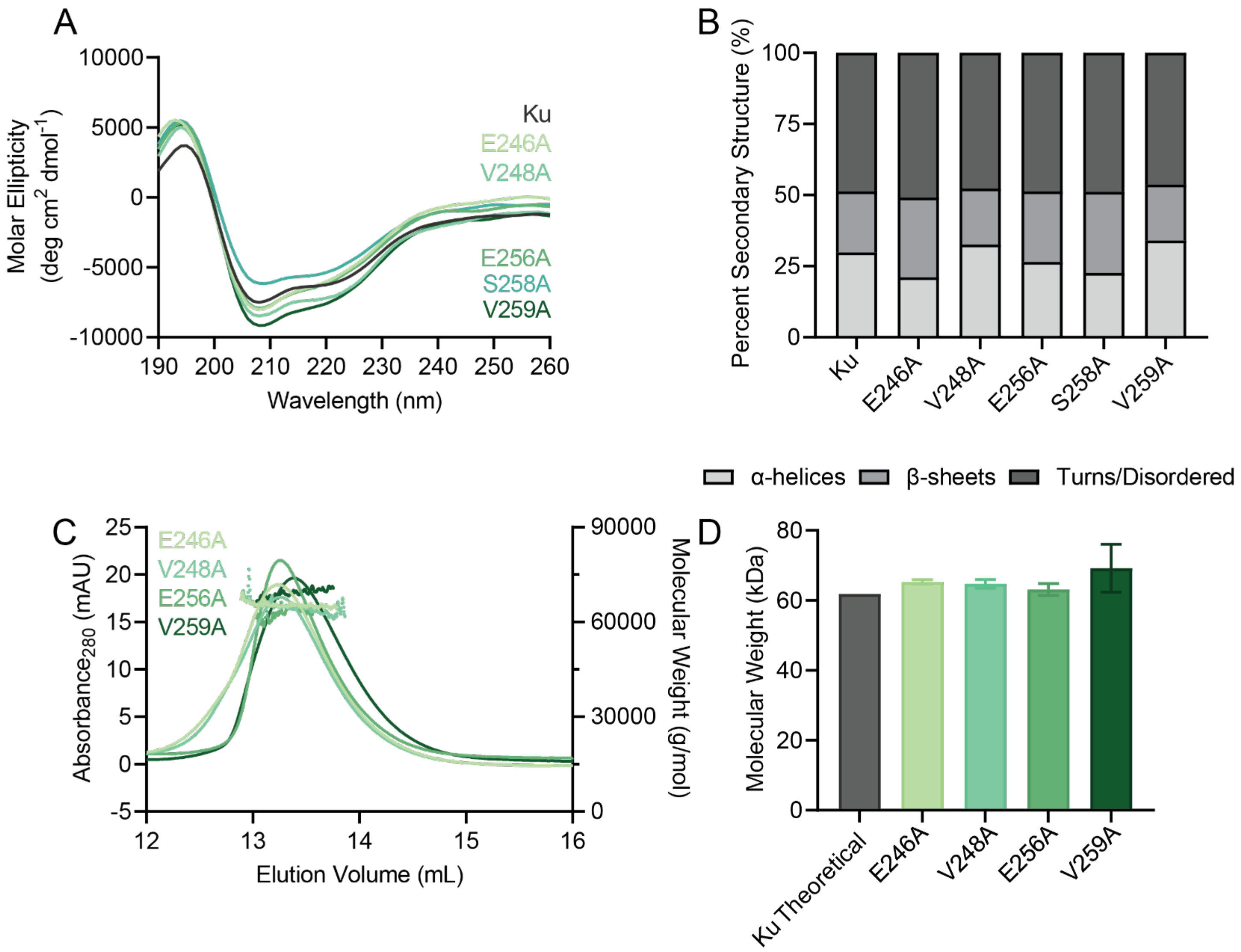


**Supplemental Figure 3:** Ku minimal C-terminal region mutants maintain wild-type Ku secondary structure. (A) Circular Dichroism Spectroscopy curves for Ku mutants compared to wild type Ku. 0.1 mg/mL samples were scanned from 260 nm to 190 nm to observe if mutations changed secondary structure. (B) Percent secondary structure for Ku mutants compared to wild-type Ku. (C) SEC-MALS data for 50 uM of Ku mutants. Elution volume is represented by the elution peaks, while molar mass is denoted as the line of best fit. (D) Molecular weight of Ku mutants compared to the theoretical molecular weight of wild-type Ku. Data are plotted as the mean ± 95% confidence interval.

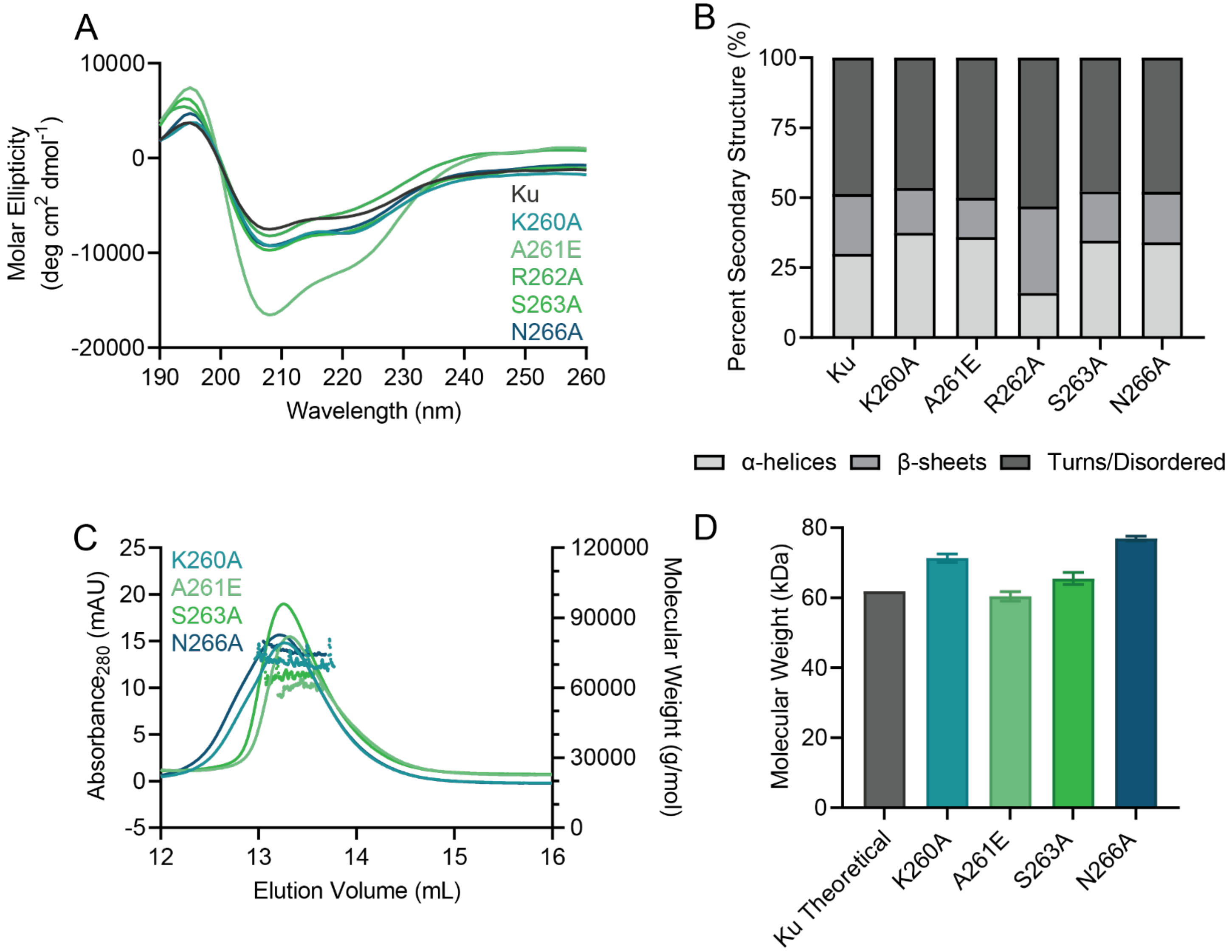


**Supplemental Figure 4:** Ku extended C-terminal region mutants maintain wild-type Ku secondary structure. (A) Circular Dichroism Spectroscopy curves for Ku mutants compared to wild-type Ku. 0.1 mg/mL samples were scanned from 260 nm to 190 nm to observe if mutations changed secondary structure. (B) Percent secondary structure for Ku mutants compared to wild-type Ku. (C) SEC-MALS data for 50 uM of Ku mutants. Elution volume is represented by the elution peaks, while molar mass is denoted as the line of best fit. (D) Molecular weight of Ku mutants compared to the theoretical molecular weight of wild-type Ku. Data are plotted as the mean ± 95% confidence interval.

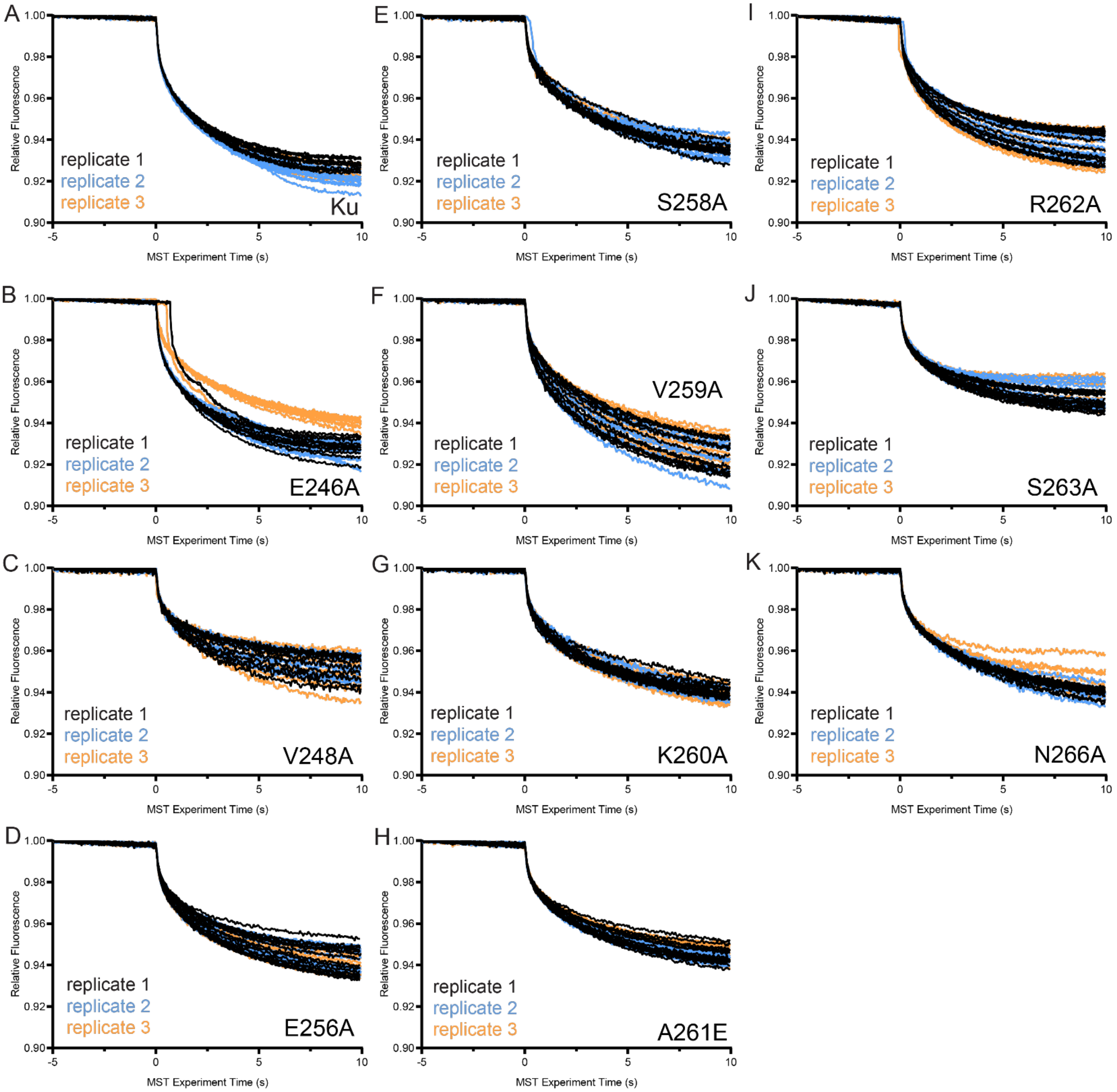


**Supplemental Figure 5:** Replicate microscale theromphoresis measurements of LigD interacting with (A) Ku and Ku mutants (B) E246A, (C) V248A, (D) E256A, (E) S258A, (F) V259A, (G) K260A, (H) A261E, (I) R262A, (J) S263A and (K) N266A. Measurements were taken over 10 seconds, n=3 technical replicates; experiments contained 12 reactions each, with the exception of E256A, A261E, R262A and S263A, which contained 16 reactions each.

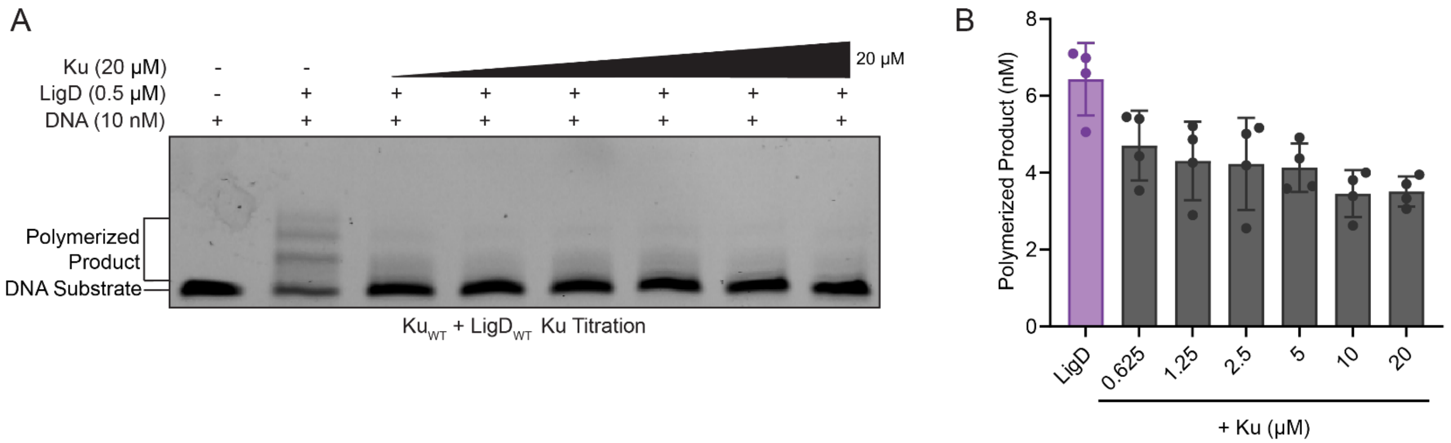


**Supplemental Figure 6:** Ku attenuation of LigD polymerase activity is not dependent on Ku concentration. (A) Representative gel image for Ku titration in templated addition assays. Reactions were run at 37˚C for a total reaction time of 5 minutes prior to loading on a 20% denaturing-PAGE gel. Electrophoresis was conducted at 200 V for 2 hours in 1X TBE running buffer. Products were visualized using the Amersham Typhoon imager (GE Healthcare). (B) Quantified polymerized product at each Ku concentration. 0.5 μM of LigD with 20, 10, 5, 2.5, 1.25, 0.625 and 0 μM of Ku. No significant difference was observed for any of the templated addition assays at various Ku concentrations.

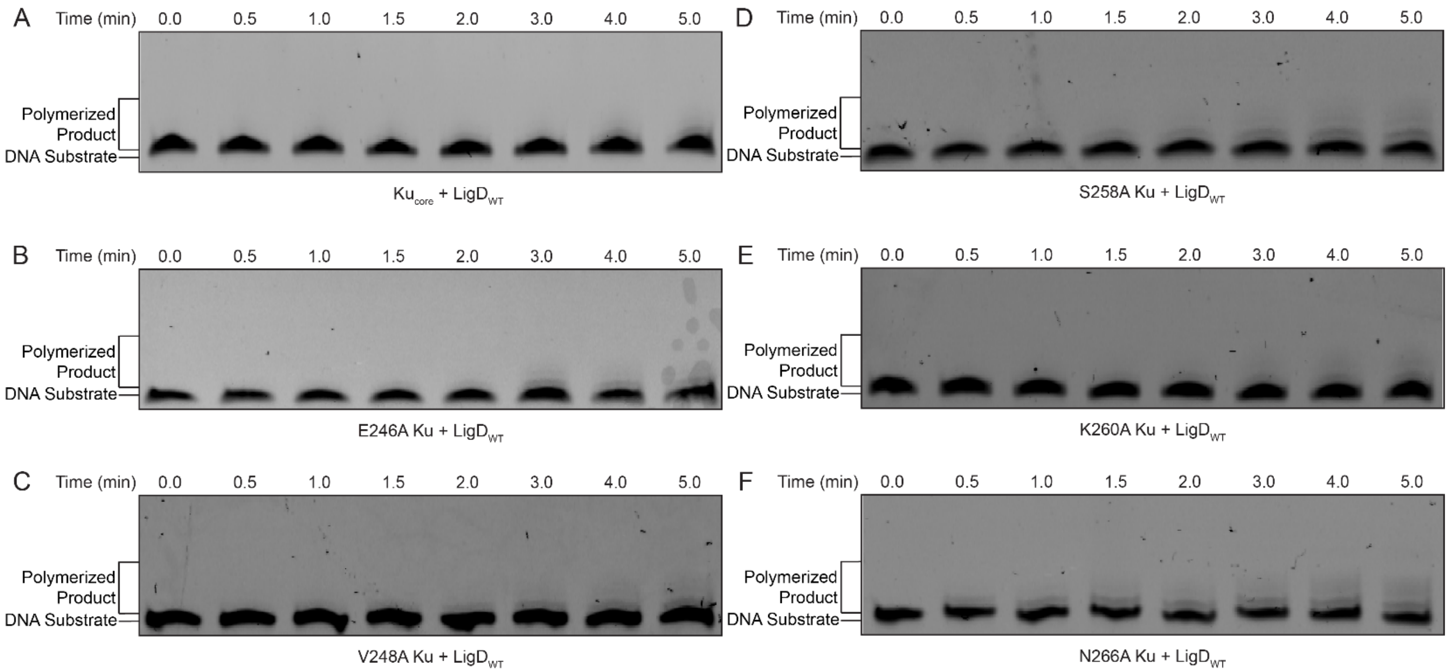


**Supplemental Figure 7:** Representative gel images for templated addition assays of Ku point mutants shown in figure 5 for LigD with (A) $Ku_{core}$, (B) Ku E246A, (C) Ku V248A, (D) Ku S258A, (E) Ku K260A and (F) Ku N266A. Reactions were run at 37˚C for a total reaction time of 5 minutes prior to loading on a 20% denaturing-PAGE gel. Electrophoresis was conducted at 200 V for 2 hours in 1X TBE running buffer. Products were visualized using the Amersham Typhoon imager (GE Healthcare).

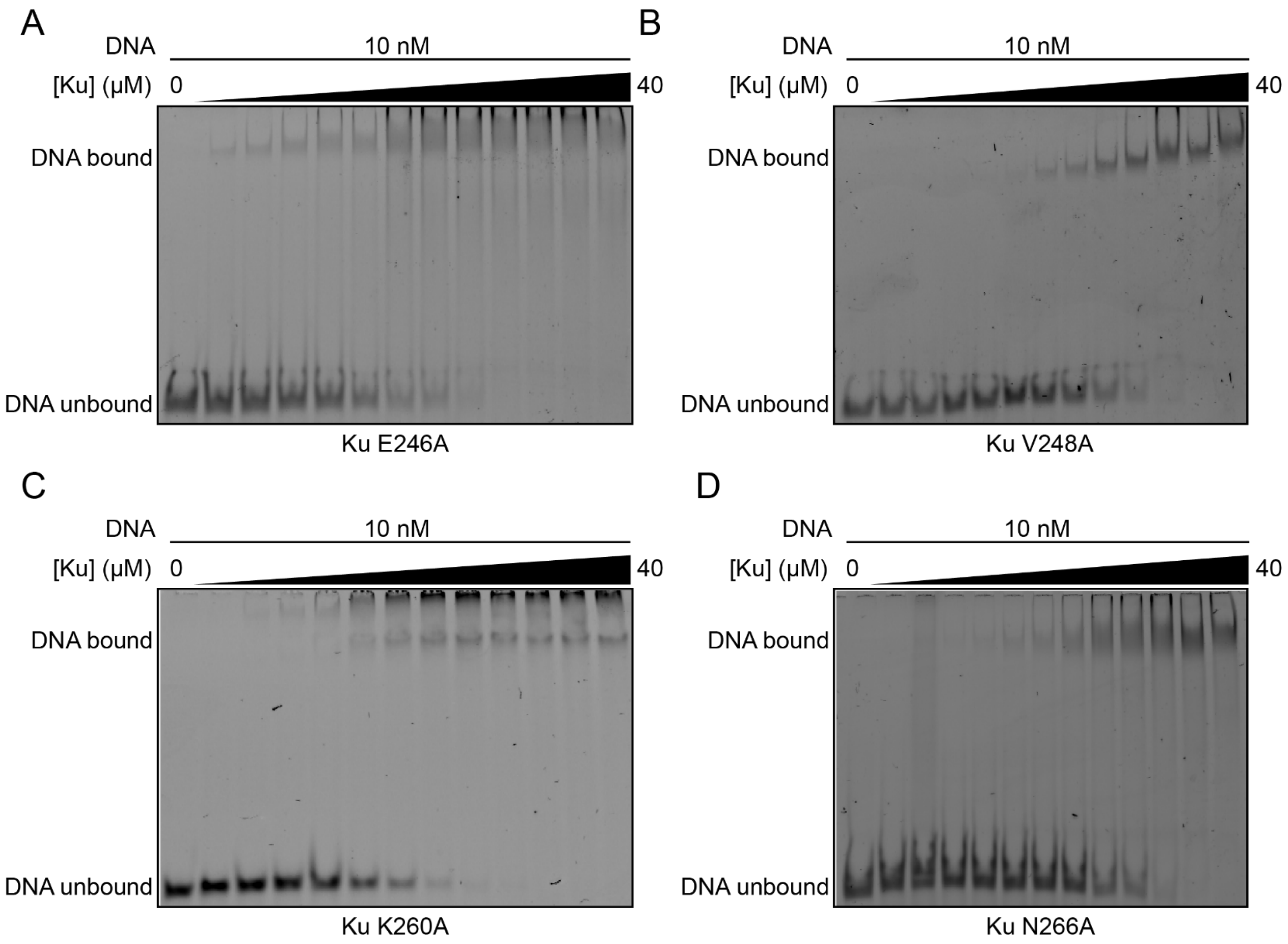


**Supplemental Figure 8:** Representative gel images for DNA binding assays of Ku point mutants shown in figure 6 for (A) Ku E246A, (B) Ku V248A, (C) Ku K260A and (D) Ku N266A. Samples were incubated at 30˚C for 20 minutes prior to loading on an 8% native-PAGE gel. Electrophoresis was conducted at 180 V for 40 minutes in 0.5X TBE running buffer. Products were visualized using the Amersham Typhoon imager (GE Healthcare).

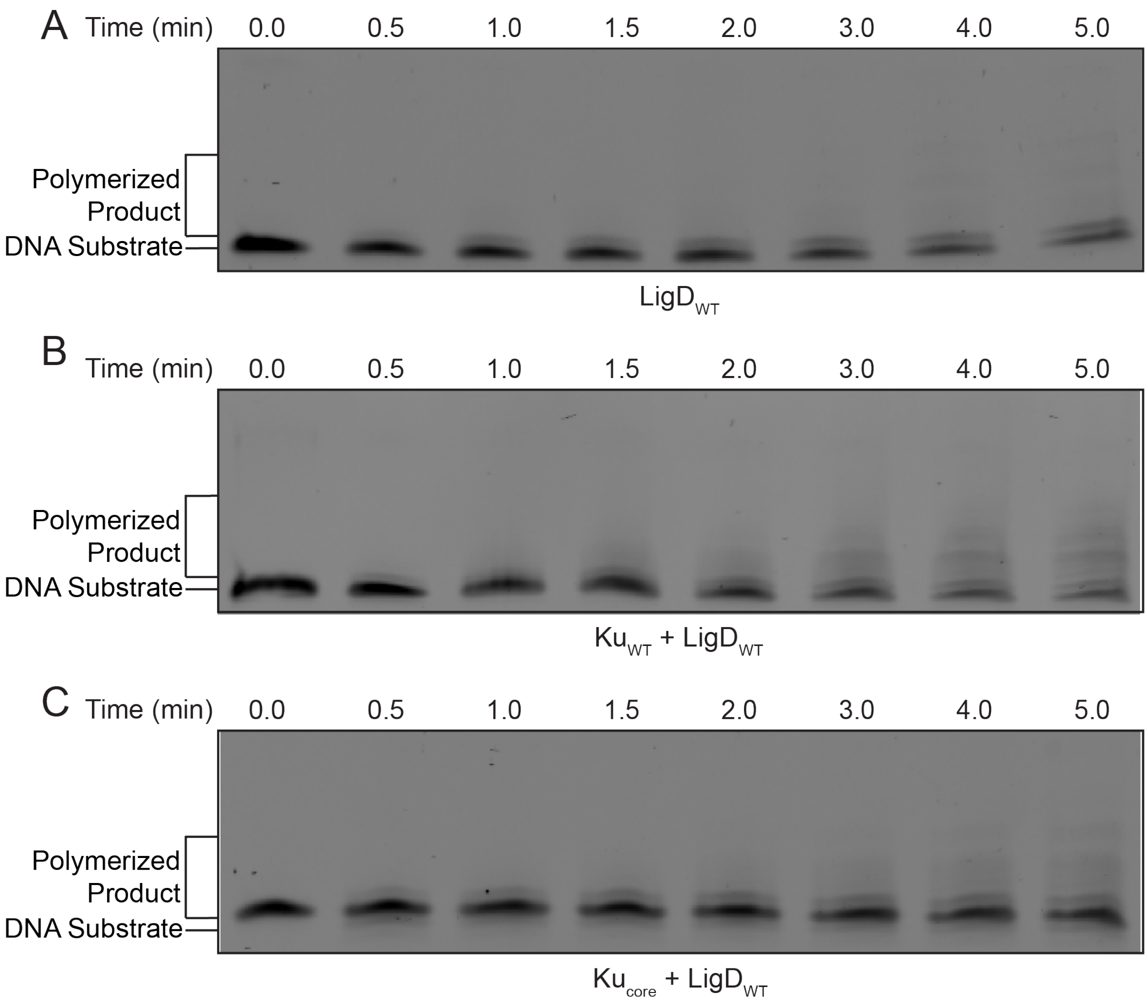


**Supplemental Figure 9:** Representative gel images for *P. aeruginosa* LigD Templated Addition Assays with Ku shown in figure 7. Representative gels for (A) LigD, (B) +Ku and (C) +$Ku_{core}$. Reactions were run at 37˚C for a total reaction time of 5 minutes prior to loading on a 20% denaturing-PAGE gel. Electrophoresis was conducted at 200 V for 2 hours in 1X TBE running buffer. Products were visualized using the Amersham Typhoon imager (GE Healthcare).

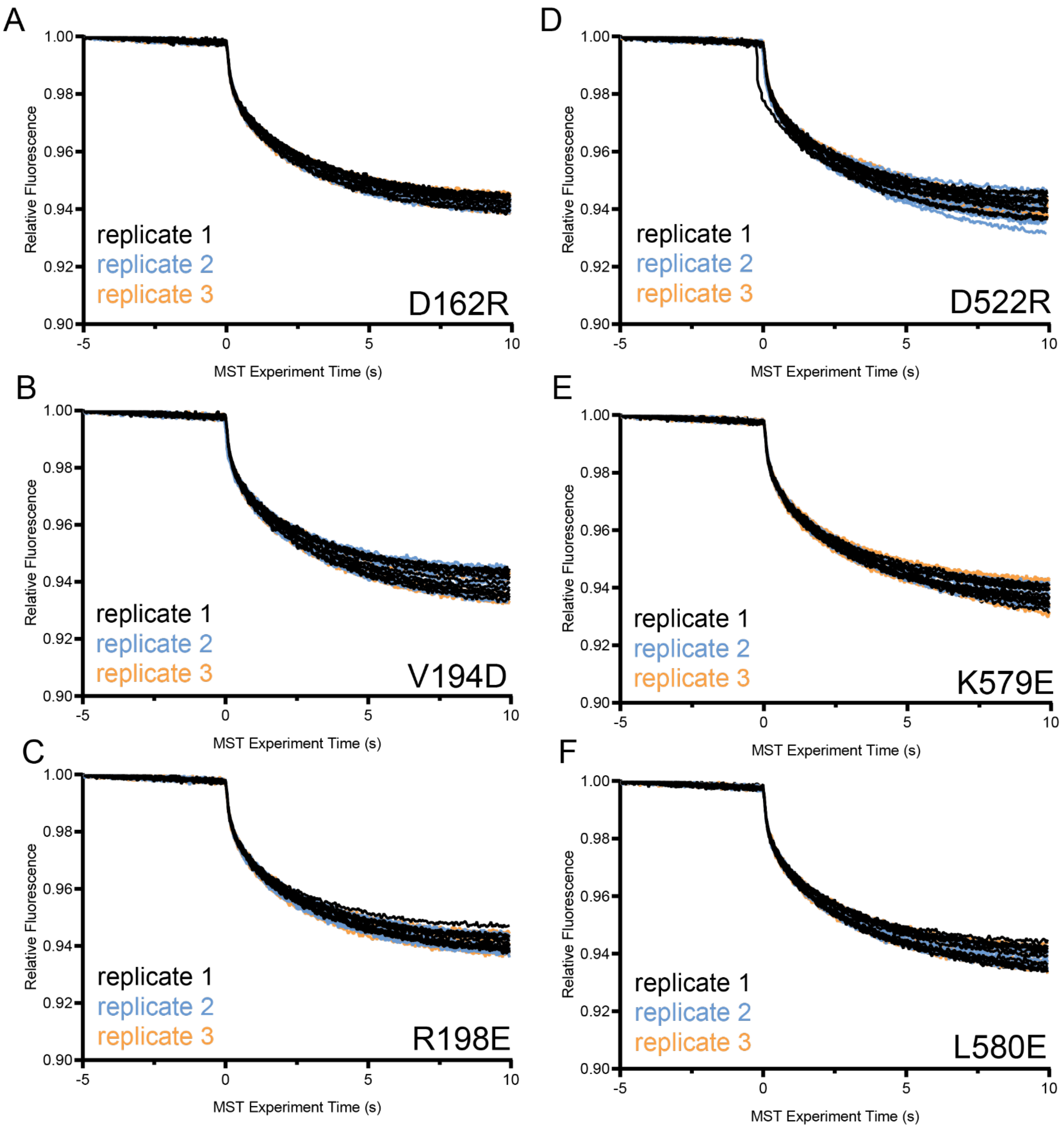


**Supplemental Figure 10:** Replicate microscale thermophoresis measurements of Ku interacting with LigD mutants (A) D162R, (B) V194D, (C) R198E, (D) D522R, (E) K579E and (F) L580E from figures 8 and 10 . Measurements were taken over 10 seconds, with n=3 technical replicates; experiments contained 12 reactions each.

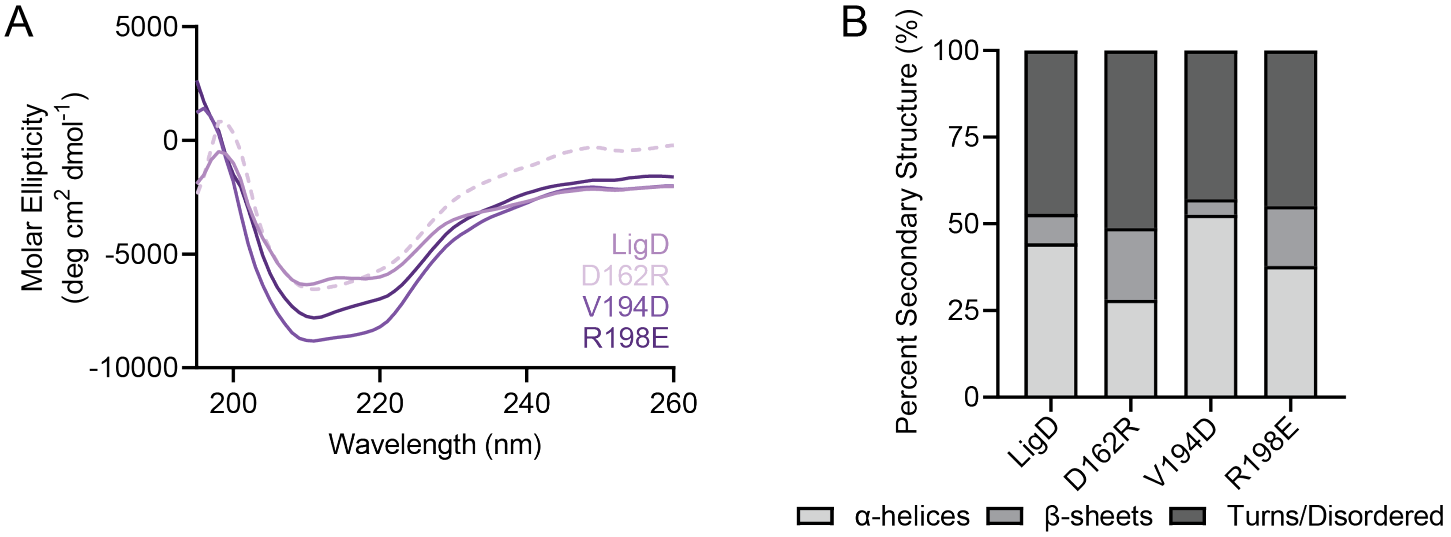


**Supplementary Figure 11:** LigD polymerase domain mutants maintain wild-type LigD secondary structure. (A) Circular Dichroism Spectroscopy curves for LigD mutants compared to wild type LigD. 0.1 mg/mL samples were scanned from 260 nm to 195 nm to observe if mutations changed secondary structure. (B) Percent secondary structure for LigD mutants compared to wild-type LigD.

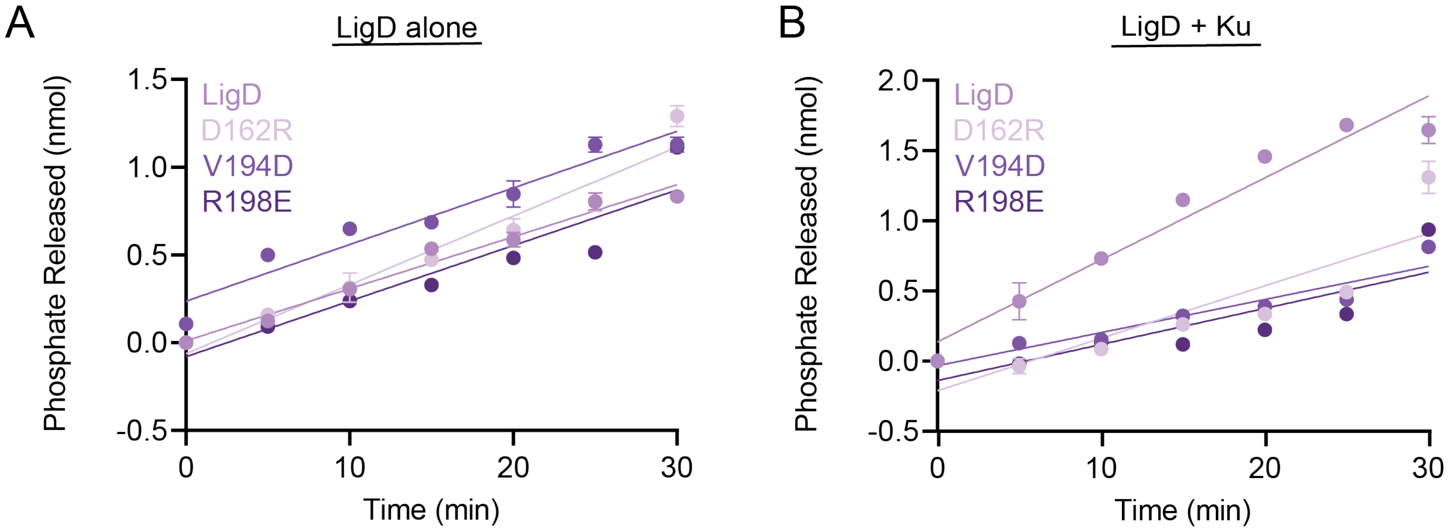


**Supplemental Figure 12:** LigD polymerase domain mutants ligase activity in the (A) absence and (B) presence of Ku for data in figure 8. Data plotted is phosphate released (nmol) vs. time (min) curves for the Biomol Green assay to assess linear rate of reaction. n=3 technical replicates. Data plotted are the mean ± standard error measure. LigD dataset is replotted here from (1) for direct comparison of data.

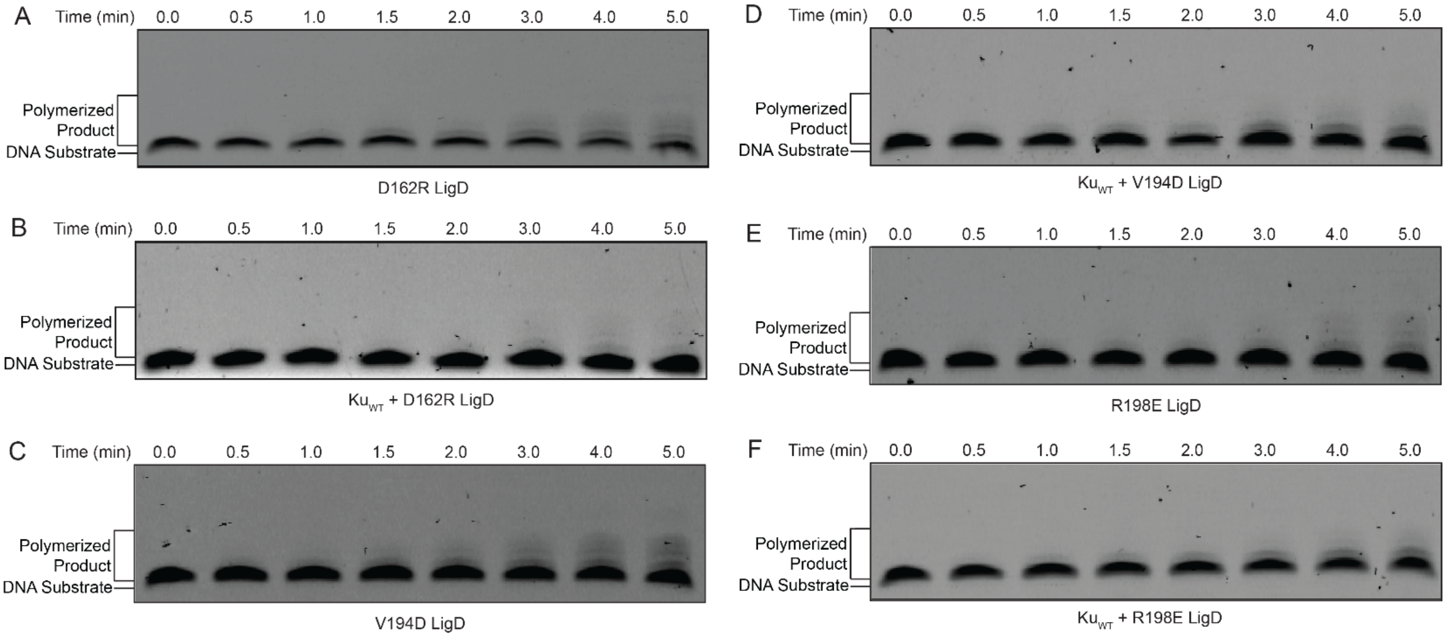


**Supplemental Figure 13:** Representative gel images for Templated Addition Assays of LigD polymerase domain point mutants shown in figure 8 for (A) LigD D162R, (B) LigD D162R + Ku, (C) LigD V194D, (D) LigD V194D + Ku, (E) LigD R198E and (F) LigD R198E + Ku. Reactions were run at 37˚C for a total reaction time of 5 minutes prior to loading on a 20% denaturing-PAGE gel. Electrophoresis was conducted at 200 V for 2 hours in 1X TBE running buffer. Products were visualized using the Amersham Typhoon imager (GE Healthcare).

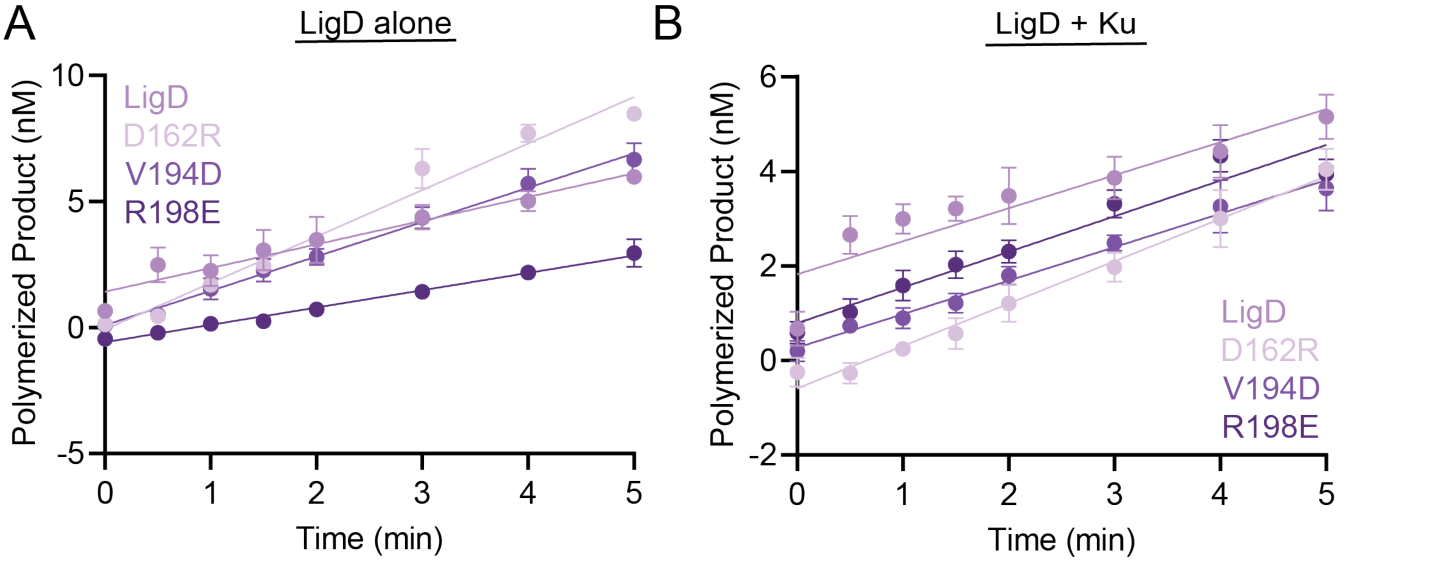


**Supplemental Figure 14:** LigD polymerase domain mutants polymerase activity in the (A) absence and (B) presence of Ku for data in figure 8. Data plotted are product polymerized (nM) vs. time (min) curves for the 18-nucleotide overhang templated addition assay to assess the linear rate of reaction. n=3 technical replicates. Data plotted are the mean ± standard error measure. LigD dataset is replotted here from Figure 4 for direct comparison of data.

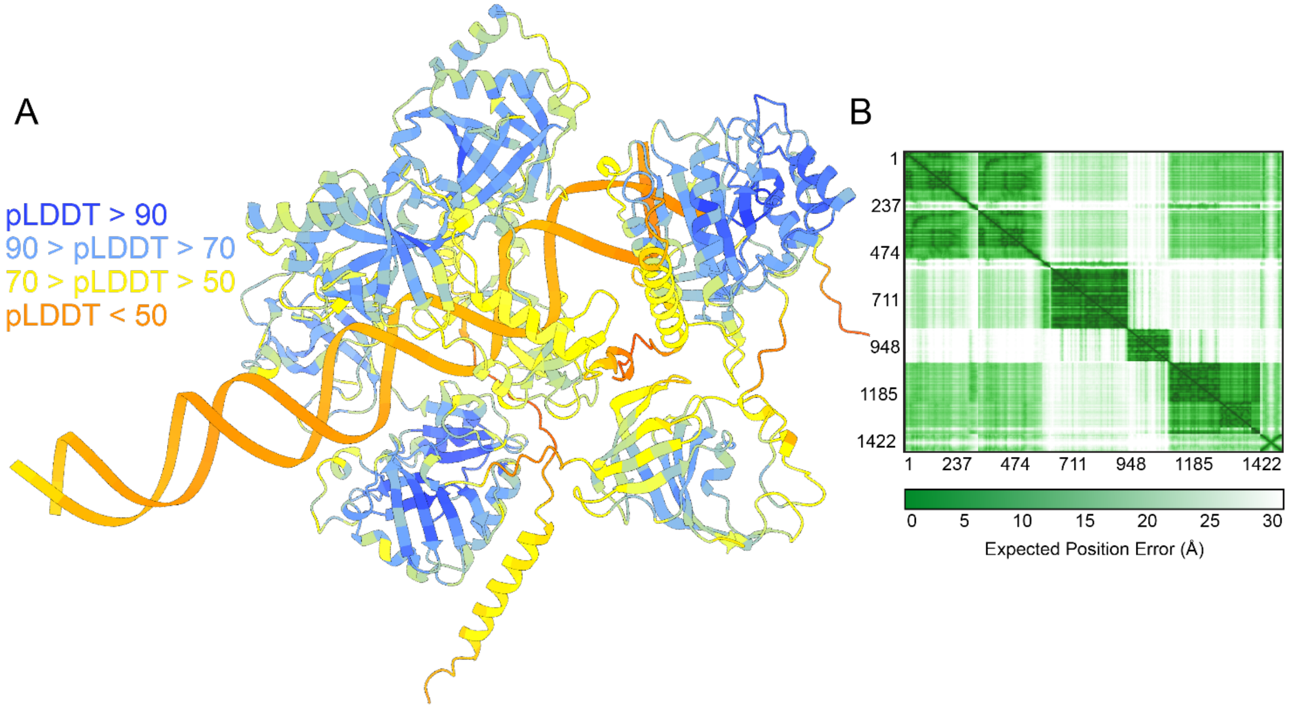


**Supplemental Figure 15:** (A) AlphaFold3 model of Ku-LigD-DNA complex colored by pLDDT in ChimeraX (version 1.9). (B) Predicted error alignment (PAE) for the AlphaFold3 model. AlphaFold3 model seen in figure 9.

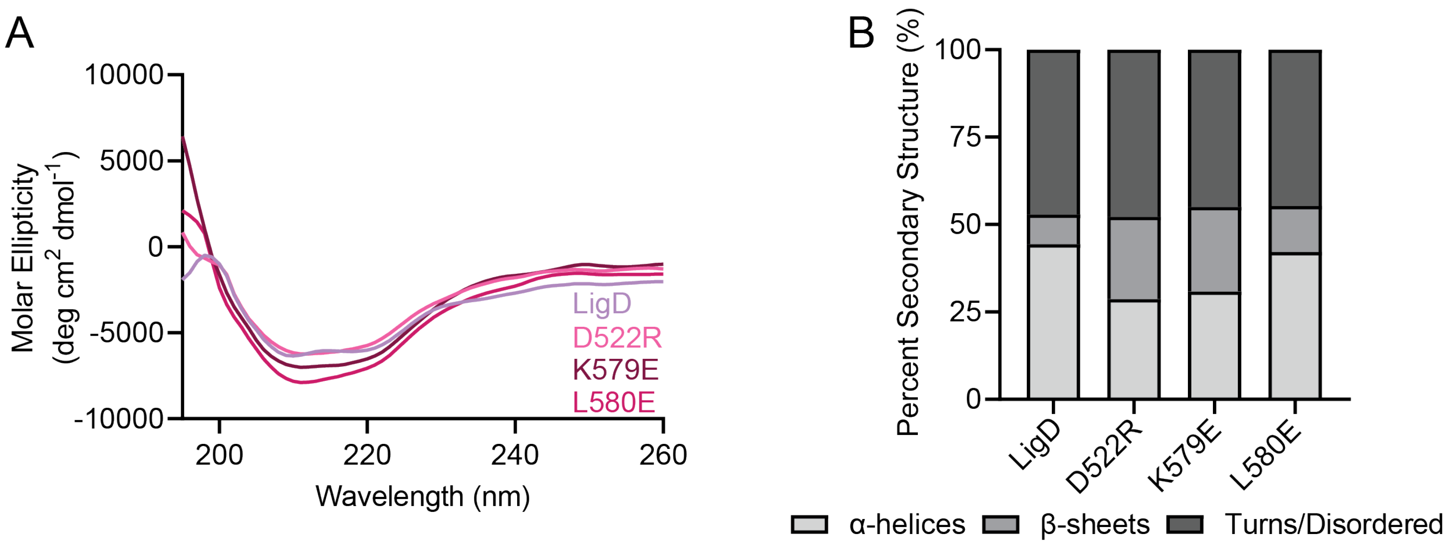


**Supplementary Figure 16:** LigD ligase domain mutants maintain wild-type LigD secondary structure. (A) Circular Dichroism Spectroscopy curves for LigD mutants compared to wild type LigD. 0.1 mg/mL samples were scanned from 260 nm to 195 nm to observe if mutations changed secondary structure. (B) Percent secondary structure for LigD mutants compared to wild-type LigD.

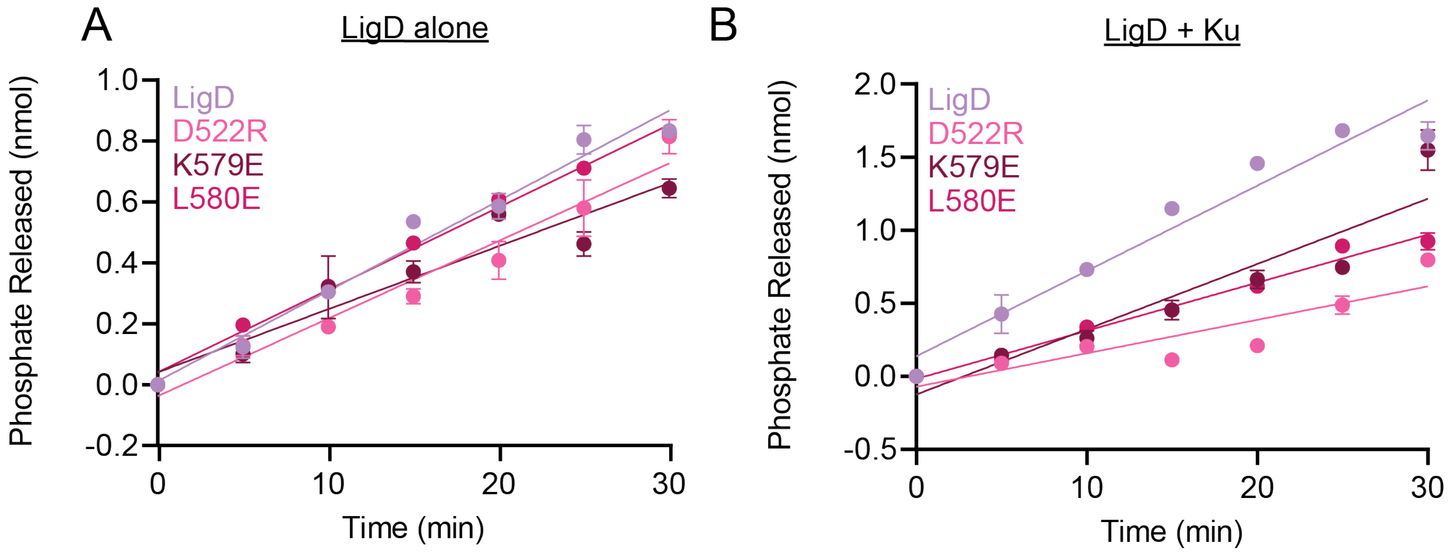


**Supplemental Figure 17:** LigD ligase domain mutants ligase activity in the (A) absence and (B) presence of Ku for data in figure 10. Data plotted are phosphate released (nmol) vs. time (min) curves for the Biomol Green assay to assess the linear rate of reaction. n=3 technical replicates. Data plotted are the mean ± standard error measure. LigD dataset is replotted here from (6) for direct comparison of data.

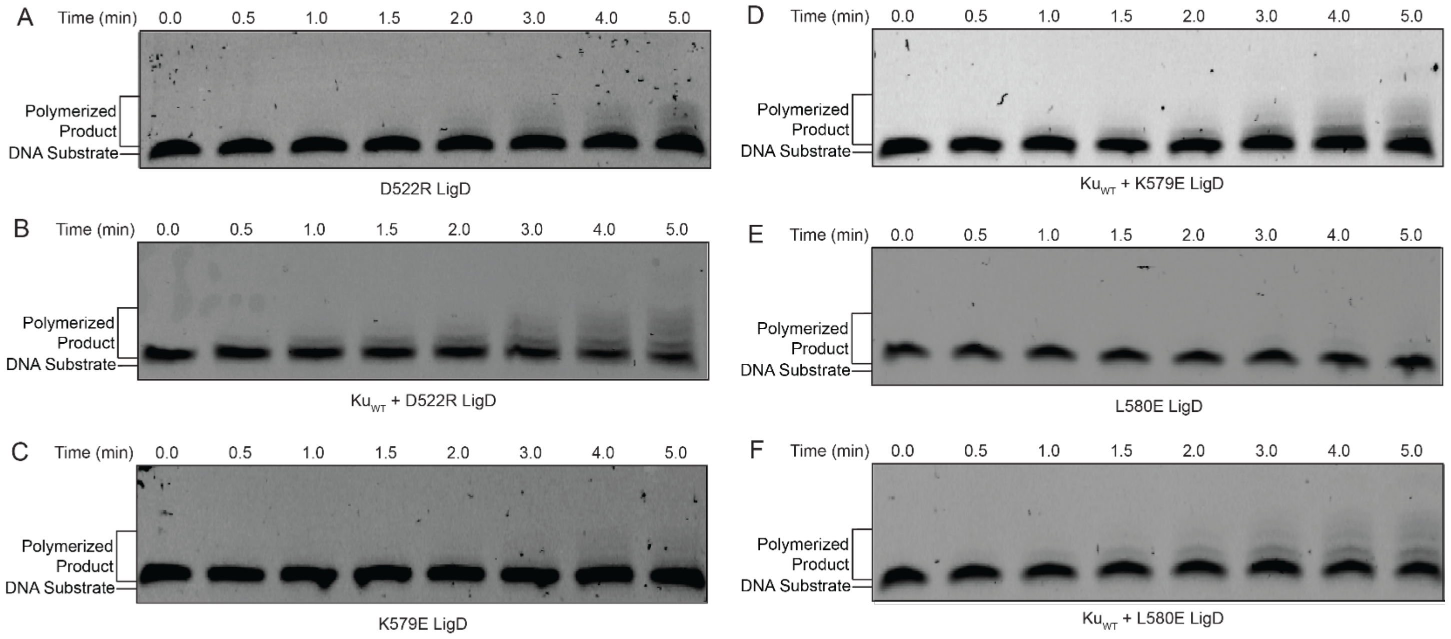


**Supplemental Figure 18:** Representative gel images for Templated Addition Assays of LigD ligase domain point mutants shown in figure 10 for LigD binding (A) LigD D522R, (B) LigD D522R + Ku, (C) LigD K579E, (D) LigD K579E + Ku, (E) LigD L580E and (F) LigD L580E + Ku. Reactions were run at 37˚C for a total reaction time of 5 minutes prior to loading on a 20% denaturing-PAGE gel. Electrophoresis was conducted at 200 V for 2 hours in 1X TBE running buffer. Products were visualized using the Amersham Typhoon imager (GE Healthcare).

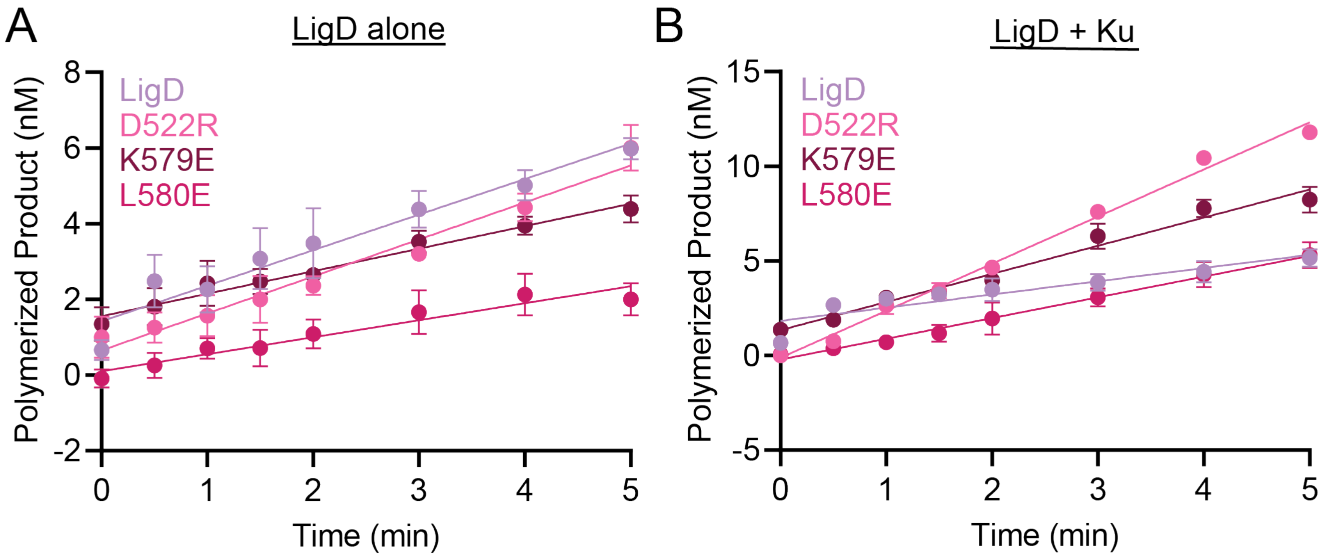


**Supplemental Figure 19:** LigD ligase domain mutants polymerase activity in the (A) absence and (B) presence of Ku for data in figure 10. Data plotted are product polymerized (nM) vs. time (min) curves for the 18-nucleotide overhang templated addition assay to assess the linear rate of reaction.  n=3 technical replicates. Data plotted are the mean ± standard error measure. LigD dataset is replotted here from Figure 4 for direct comparison of data.

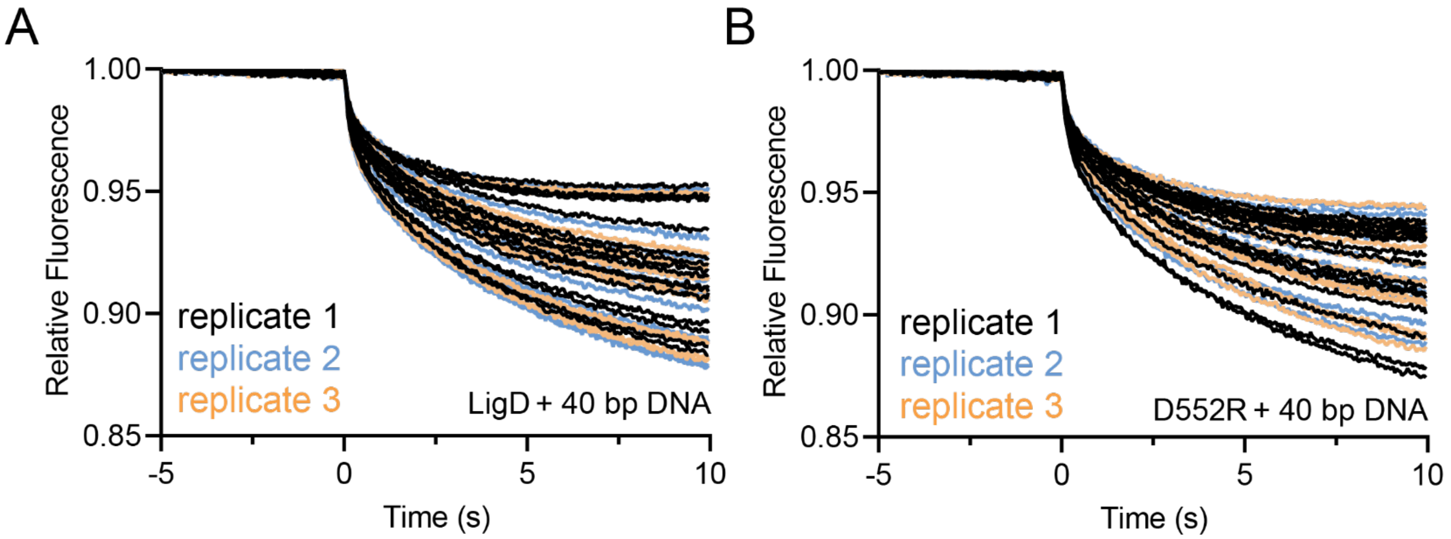


**Supplemental Figure 20:** Replicate microscale therомphoresis measurements of a Ku-DNA complex interacting with (A) LigD and (B) LigD mutant  D522R for data in figure 10. Measurements were taken over 10 seconds, with n=3 technical replicates; experiments contained 16 reactions each.
